# Discovery and evaluation of cadmium-adapted *Daphnia pulex* genotypes in a region of historical mining reveals adaptation protects the germline from cadmium-induced mutations

**DOI:** 10.1101/2025.11.11.687873

**Authors:** Nathan Keith, Stephen P. Glaholt, Craig E. Jackson, Kim Young, Karel De Schamphelaere, John K. Colbourne, Joseph R. Shaw

## Abstract

Exposure to chemical pollutants can alter the rate, and genome-wide distribution of germline mutations. However, studies measuring the effect of chemical exposure on mutation rate and spectra have not considered the ecological and evolutionary backgrounds of studied genotypes, which could influence the rates and patterns of germline mutations in altered environments, e.g., chemical pollution. Utilizing a study of natural *Daphnia pulex* populations, we conducted a comprehensive experiment to test our hypothesis that adaptation to chemical pollution also protects the germline from mutagenesis. We, 1) identified *Daphnia pulex* populations that have adapted to live in mining-devastated regions by increasing their cadmium tolerance. 2) We completed a mutation-accumulation (MA) experiment with an adapted genotype to measure the germline mutation rate in both control conditions and an environmentally relevant cadmium concentration. 3) We compared these MA experiment results to a previously reported, identically designed MA experiment with a nonadapted genotype. We report that patterns of cadmium-induced mutagenesis in the adapted genotype were reversed compared to our previous observations in the nonadapted genotype. Cadmium exposure altered the single nucleotide mutation (SNM) rate in the same genome regions in adapted and nonadapted genotypes, but the rates were changed in opposite directions. Cadmium also altered specific SNM classes in these genotypes in opposite directions. The reversal of mutational trends in the adapted genotype suggests protection against cadmium genotoxicity. We further demonstrate that adapted populations have elevated gene copy-number and expression levels of metallothionein, the protein that protects against cadmium toxicity by binding to cadmium irreversibly.

## Introduction

Organisms can adapt to live in extreme environments, including those that are heavily polluted by chemicals (Reid, et al. 2016). Understanding the coping mechanisms that allow for population survival in the presence of toxic concentrations of chemicals is important because of the increasing rate of pollution over the past century (Landrigan, et al. 2018). The ultimate source of the genetic variation that allows populations to adapt are germline mutations (Denver, et al. 2004; Lynch 2010; Sung, Tucker, et al. 2012a; Schrider, et al. 2013; Sung, et al. 2015; Keith, et al. 2016).

Selection acts on this variation and promotes genotypes offering a fitness advantage in a given environment. If a selective force is strong, as can be the case for environments contaminated with chemicals, and if certain population genetic parameters are met within a population, affected genotypes can become highly specialized. Although germline mutations can be beneficial to fitness, the majority have a neutral or mildly deleterious impact on fitness (Lynch, et al. 1999), and this would seem especially true for these specialized, environmentally matched, adapted genotypes.

Several studies have documented physiological adaptation in response to heavily polluted environments. Reid et al. (2016) applied a population genomics approach to identify many genomic loci of reduced variability in four populations of killifish, *F. heteroclitus*, that independently adapted to survive in highly polluted environments that are toxic to non-adapted populations (Reid, et al. 2016). They conclude that selection constrains standing variation in these populations, often in similar genomic regions, to produce the adaptive phenotype. Many have used common garden or reciprocal transplant studies to adaptation to polluted environments in multicellular organisms (Bergelson and Purrington 1996; Guedes, et al. 2006; Salice, et al. 2010; Jansen, et al. 2011; Dutilleul, et al. 2017), but the mutational processes in these adapted genotypes were not investigated.

Mutation accumulation (MA) experiments provide a powerful, yet underutilized, opportunity for measuring the environmental influences on the rate and spectrum of germline mutation in adapted genotypes. MA experiments remove natural selection via strict genetic bottlenecks after organismal reproduction each generation (Halligan and Keightley 2009; Sung, Ackerman, et al. 2012; Keith, et al. 2016; Sung, et al. 2016). The removal of natural selection preserves all germline mutations except the extreme minority that cause immediate lethality or sterility. When propagated for thousands of generations and subjected to whole-genome sequencing, MA experiments provide precise estimates of de novo germline mutational processes. Tightly controlled MA experiments that include chemically adapted genotypes maintained in both presence and absence of the adaptive environment, therefore, provide a unique opportunity for understanding mutational processes in adapted genotypes.

We recently reported that environmentally relevant concentrations of cadmium can drastically alter the rate and spectrum of germline mutation (Keith, et al. 2021). After analysis of a large-scale *Daphnia pulex* mutation-accumulation (MA) experiment in control and cadmium exposure, we showed that cadmium exposure changes the mutation rate in multiple genome regions, while also changing the rates of multiple classes of single nucleotide mutations (SNMs). Such environmentally driven changes in mutation rates and spectra, could be detrimental to populations adapted to polluted environments.

Utilizing *D. pulex*, we conducted a comprehensive experiment to test our overarching hypothesis that adaptation to chemical pollution also protects the germline from cadmium-induced mutagenesis. We, 1) identified *Daphnia pulex* populations that have adapted to live in mining-devastated regions by increasing their tolerance to cadmium. 2) We completed a large-scale MA experiment with an adapted genotype to measure the rate of germline mutation in both control conditions and an environmentally relevant concentration of cadmium. 3) We compared these MA experiment results to a previously reported and identically designed MA experiment with a non-adapted genotype.

## Results

### Discovery of cadmium-adapted *Daphnia pulex* genotypes in Sudbury, Ontario

Our study focused on representative genotypes from 13 lakes in Ontario, Canada (Fig. 1A, Table S1), including seven highly acidified, metal-contaminated lakes near Sudbury, a center of nickel mining and smelting since the 1800s (Dixit, et al. 1992; Tropea, et al. 2010; Schindler and Kamber 2013) and six lakes near Dorset, 200 km to the Southeast, which do not share the Sudbury lakes’ metal-contamination history. A microsatellite based phylogenetic study of *D. pulex* delimits with 100% bootstrap support a cluster containing four Sudbury populations, Group A (Shown in red in Fig. 1B). The three other Sudbury lake populations clustered with Dorset populations, Group B (Fig. 1B).

**Figure 1.**
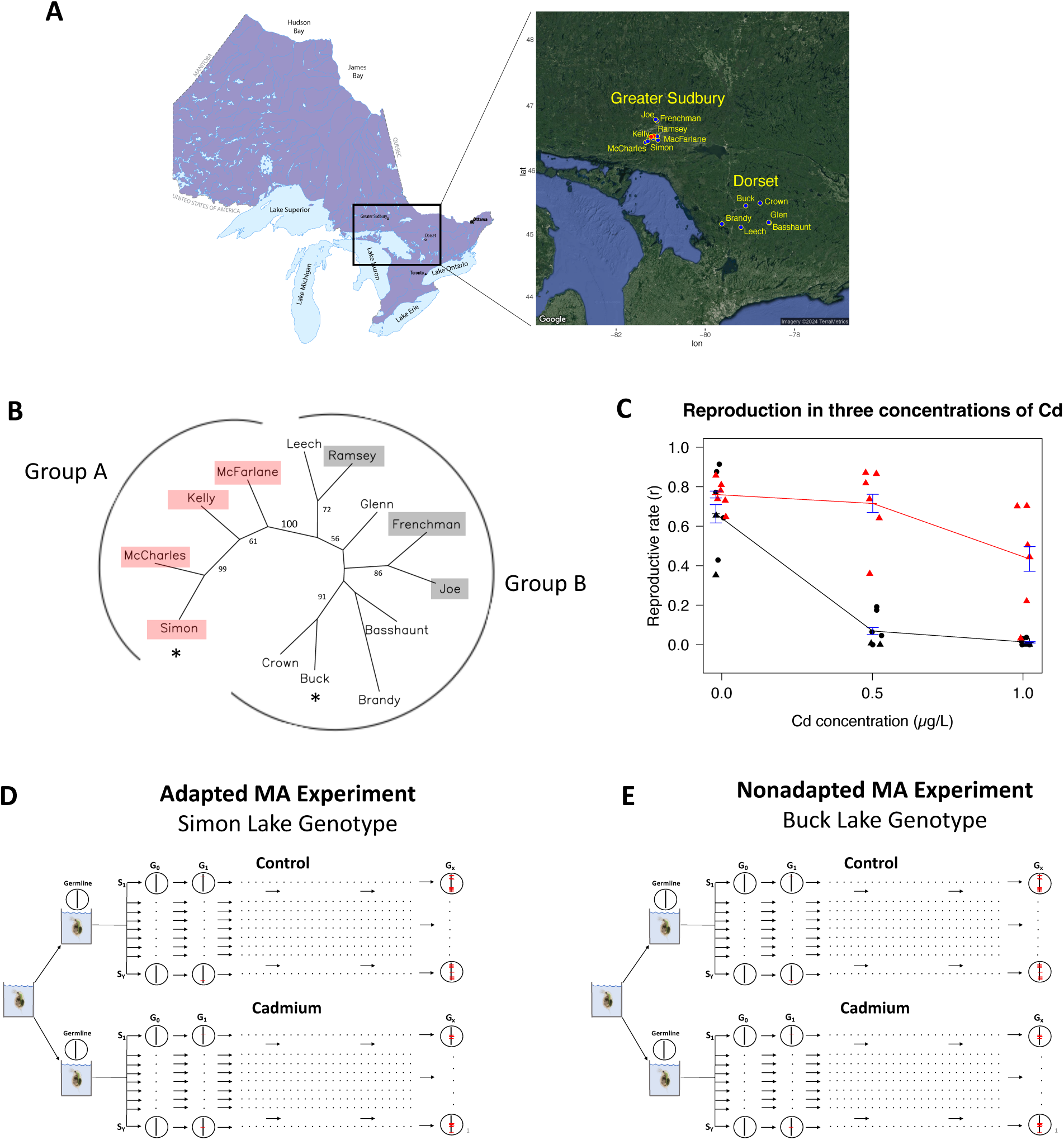
Sudbury and Dorset lake populations, cadmium toxicity and MA experimental design. (A) Latitude/longitude coordinates of lakes sampled in Sudbury, Ontario and Dorset, Ontario. Black points represent the locations of lakes, while red points represent the locations of Sudbury mines. (B) Consensus NJ tree of Cavalli-Sforza genetic distances based on 1,000 bootstraps of allele-frequency data for 26 microsatellites across 192 strains from 13 lake populations. Bootstrap values above 50% are shown. The branch separating cadmium-tolerant populations and cadmium-sensitive populations has 100% bootstrap support. Sudbury populations are shown in shaded rectangles, which are red for the adapted lake populations. (C) Reproductive rate (offspring/day) during 21-day chronic exposure to three concentrations of cadmiums. Red triangles: tolerant, Sudbury genotypes; Black triangles: sensitive Sudbury genotypes; Black circles: sensitive Dorset genotypes. (D, E) Overview of mutation-accumulation experimental design. For both adapted and non-adapted genotypes, a single female was used to initiate each experiment. Each experiment (i.e. non-adapted and adapted) were founded with genetically identical females all, descended from the original “mother”. Both genotypes were then propagated in control and chronic, cadmium exposure. The vertical lines within circles represent the germline genome. Red horizontal lines on the germline genome are representative of mutations that occur in the course of a MA experiment. “*G*”, germline. “*S*”, Subline. “*Y*”, total number of sublines. “*X*”, total number of generations.

All genotypes were acclimated to clean laboratory, common garden conditions for 25 generations (Methods). Following acclimation, genotype specific reproduction rates were measured using 21-day reproductive fitness assays in cadmium, as well as, nickel, and arsenic (USA 2002), which are also observed in elevated concentrations in Sudbury area lakes (Nriagu, et al. 1982; Lock, et al. 2016). Although exposure to nickel and arsenic significantly reduced fitness in all groups, Group A genotypes from Sudbury were tolerant to cadmium (Figure 1B, Figure 1C, Figure S1, Tables S2, S3). At 0.5 µg cadmium/L, Group A strains produced offspring at 94% the rate of control conditions, while reproduction in the sensitive genotypes, Group B, was only 9% of control conditions (*P* = 2.5 x 10^-3^, Wilcoxon, Fig. 1C).

The cadmium specific tolerance of Group A was also observed when the cadmium concentration was doubled. At 1 µg cadmium/L reproductive rates in Groups B were reduced to 2% of control conditions, while Group A was 56% relative to control conditions (*P* = 4.7 x 10^-3^, Wilcoxon, Fig. 1C). The results of our fitness assays due to multi-generation culture in clean, common garden conditions eliminate the possibility of physiological acclimation to cadmium, indicate that the observed tolerance is heritable and genome-encoded (i.e., adaptation) because these cultures were held in clean (no cadmium), common garden conditions for over 10 generations prior to toxicity testing.

Like most *Daphnia* species, *D. pulex* and *D. pulicaria* are members of a larger species complex that actively engage in hybridization and introgression (Colbourne et al., 1998, Crease et al. 2012) as evident by the ∼60% of the genomes of these two species that are historically homogenized via gene flow (Jackson et al., 2021). Historically, the allozyme locus *Ldh* has been used to identify these species and their hybrids via electrophoresis. *D. pulex* are characterized by homozygosity for a slow (S) *Ldh*A allele, while *D. pulicaria* are characterized by homozygosity for a fast (F) *Ldh*A allele, and hybrids are heterozygous containing both slow and fast (SF) *Ldh*A alleles (Crease et al. 2011). ADAP and Group A (Figure 1B) genotypes were phenotyped as slow-fast (SF) *Ldh*, indicating hybridity, while NONA was phenotyped as fast-fast (FF) *Ldh*. For experimental ease, the chosen isolates for this study reproduce by obligate asexuality, ensuring clonality over multiple generations. Tucker et al. showed that obligate asexuality in *D. pulex* is caused by specific chromosome 8 and 9 haplotypes introgressed from *D. pulicaria* and maintained as heterozygous alongside the *D. pulex* chromosome 8 and 9 haplotypes (2013). If this is only path to obligate asexuality in *D. pulex*, NONA must also contain introgressed haplotypes from *D.pulicaria* and exhibit hybridity on chromosomes 8 and 9.

Before initiating these studies, we wanted to explicitly eliminate the possibility that cadmium tolerance was a general feature of hybrids, i.e., through hybrid vigor (heterosis). For this, we assessed the reproductive fitness of F_1_ hybrids following cadmium exposure compared to their parental strains (i.e., “pure” *D. pulex* and *D. pulicaria,* Figure S1). We tested 17 F1 hybrids, and 5 *D. pulex* and 9 *D. pulicaria* isolates that served as the parental strains for these hybrids in control conditions and two cadmium concentrations, 2.5 and 5 μg Cd/L. While hybrid vigor was observed when comparing F_1_ hybrids to the mid-parent value for *D. pulex* and *D. pulicaria* strains in control conditions (P = 0.006, Figure S1), following cadmium exposure no significant differences were observed between hybrids and either *D. pulex* parental strains (P = 0.82 for 2.5 μg Cd/L, P = 0.43 for 5 μg Cd/L), *D. pulicaria* parental strains (P = 0.53 for 2.5 μg Cd/L, P = 0.65 for 5 μg Cd/L) or their mid-parent values (P = 0.33 for 2.5 μg Cd/L, P = 0.21 for 5 μg Cd/L) (Fig. S1). In addition, the rate of reproductive decline in response to increasing cadmium concentration was not significantly different between hybrids and the parent species (*D. pulex* parental P = 0.077, *D. pulicaria* parental P = 0.086). We conclude that the hybrid vigor observed in optimal conditions and *D.pulex* × *D. pulicaria* hybridization itself does not confer an enhanced tolerance to cadmium.

### Mutation-accumulation experiment with an adapted *D. pulex* genotype

To address our hypothesis that adaptation protects the germline from chemical-induced changes in mutational processes, we propagated an MA experiment for 1,776 generations with a cadmium-adapted genotype in both control conditions and cadmium exposure (Fig. 1D) (hereafter referred to as ADAP Control and ADAP Cd). Following 1,776 generations of MA, 12 ADAP Control sublines representing 741 total generations, and 12 ADAP Cd sublines comprising 1035 generations were subjected to whole-genome, deep-coverage sequencing (average sequencing coverage ∼25x, Supplemental Table S4). Control and cadmium exposure MA sublines were maintained in laboratory conditions as described in Keith et al. (2021). We compared the mutational patterns of ADAP Control to ADAP Cd to measure the influence of cadmium on germline mutation in this genotype. We then compared the germline mutational response to cadmium in ADAP to the results of our previously reported, identically designed MA experiment conducted in parallel with a nonadapted *D. pulex* genotype (Keith, et al. 2021) (hereafter referred to as NONA Control and NONA Cd; Fig. 1E).

Keith et al. (2021) reported that cadmium dramatically changed genome-wide patterns of mutation. Specifically, cadmium exposure changed the transition to transversion (Ts/Tv) ratio and caused an increased G/C → T/A mutation bias (i.e. the number of mutations in the G/C → T/A direction compared to the number of mutations in the T/A → G/C direction). Cadmium exposure also changed the conditional rate of mutation for multiple mutation classes. Finally, cadmium exposure dramatically changed the genome-wide distribution of mutations by changing the intergenic, genic, and exon rates of mutation. We therefore investigated these same mutational classes and changes in ADAP for comparison between the two genotypes with differing evolutionary histories. Additionally, we performed multiple physiological and reproductive rate analyses for the NONA and ADAP genotypes to better characterize the functional basis of adaptation.

### Reproductive Rate (r) Analyses

Reproductive rate (r), a quantitative measurement of *D. pulex* fitness, was measured at multiple generation timepoints (Generations 10, 25, 40) of the NONA and ADAP MA experiments. We observe opposite trends in reproductive rate for NONA and ADAP throughout the course of the experiments. In ADAP after 10 generations (generation 10), reproductive rate is significantly lower in cadmium exposure (T-test, *P* = 3.3 x 10^-5^), while reproductive rate is higher in cadmium exposure at generation 25 (T-test, *P* = 9.3 x 10^-51^) and generation 40 (T-test, *P* = 2.1 x 10^-25^) (Fig. 2A). Contrary to ADAP, in the NONA experiment cadmium exposure significantly decreases reproductive rate at generation 25 (T-test, *P* = 6.4 x 10^-5^) and generation 40 (T-test, *P* = 2.2 x 10^-4^) (Fig. 2B), while no significant difference was observed at generation 10 (T-test, *P =* 0.5).

**Figure 2.**
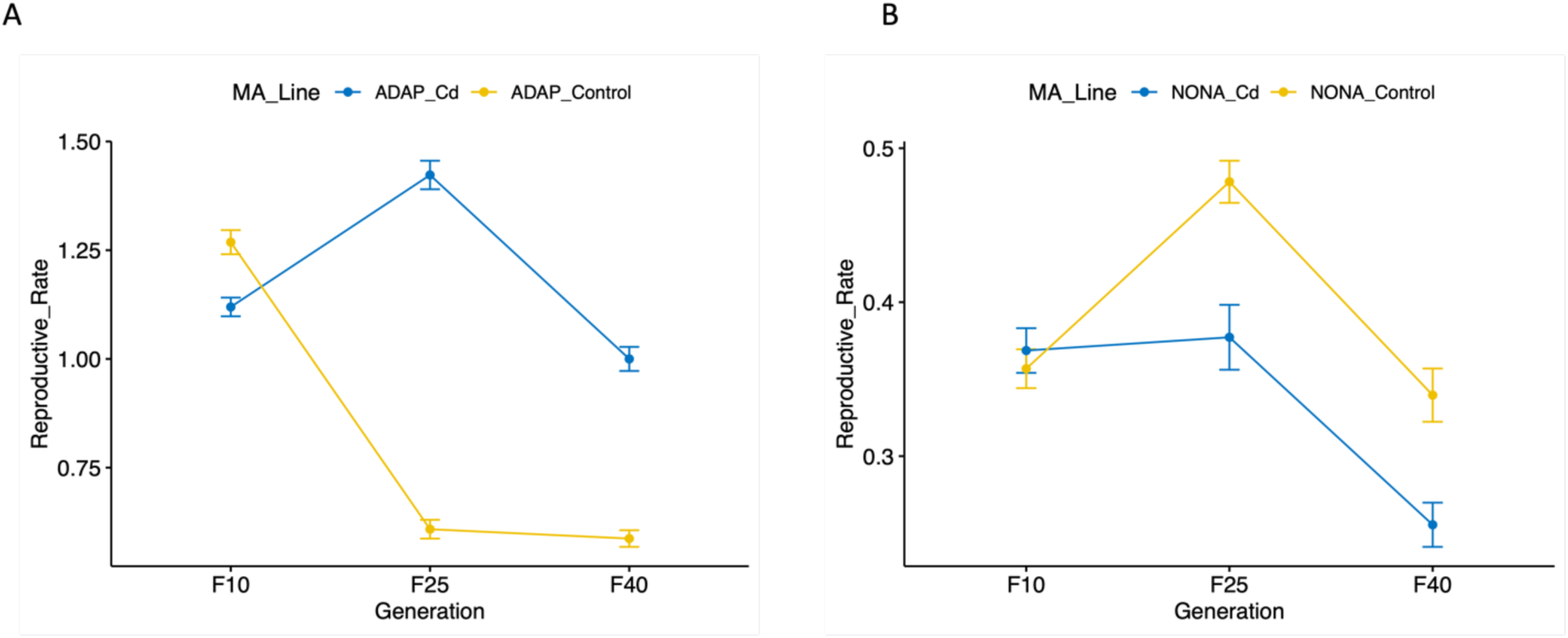
Superoxide dismutase (SOD) and Glutathione S-transferase (GST) (B) results. (A) GST activity results. Y-axis values are measured in units µmol / ml / min, normalized by the volume of input supernatant (Methods). (B) SOD activity results. Y-axis values are measured in units of percent inhibition (i.e., activity; Methods).

### Glutathione S-transferase (GST) Activity

We measured GST and SOD activity to determine if antioxidant enzymes were playing a part in adaptation to cadmium. For GST, in control conditions (0 μg Cd / L) and cadmium exposure (0.25 and 20 μg Cd / L) with both the ADAP and NONA genotypes (Materials and Methods). In ADAP, compared to control conditions 0.25 μg Cd / L did not increase GST activity (T-test, *P* = 0.19), but GST activity was significantly increased by 20 μg Cd / L exposure (T-test, *P* = 0.009, Fig. 3A). In NONA, relative to control conditions, cadmium exposure (at concentrations of both 0.25 and 20 μg Cd / L) did not significantly increase GST levels (T-test, *P* = 0.24 and 0.30 for 0.25 and 20 μg Cd / L, respectively). However, at 20 μg Cd / L GST is significantly higher in ADAP compared to NONA (T-test, *P* = 0.03, Fig. 3A).

**Figure 3.**
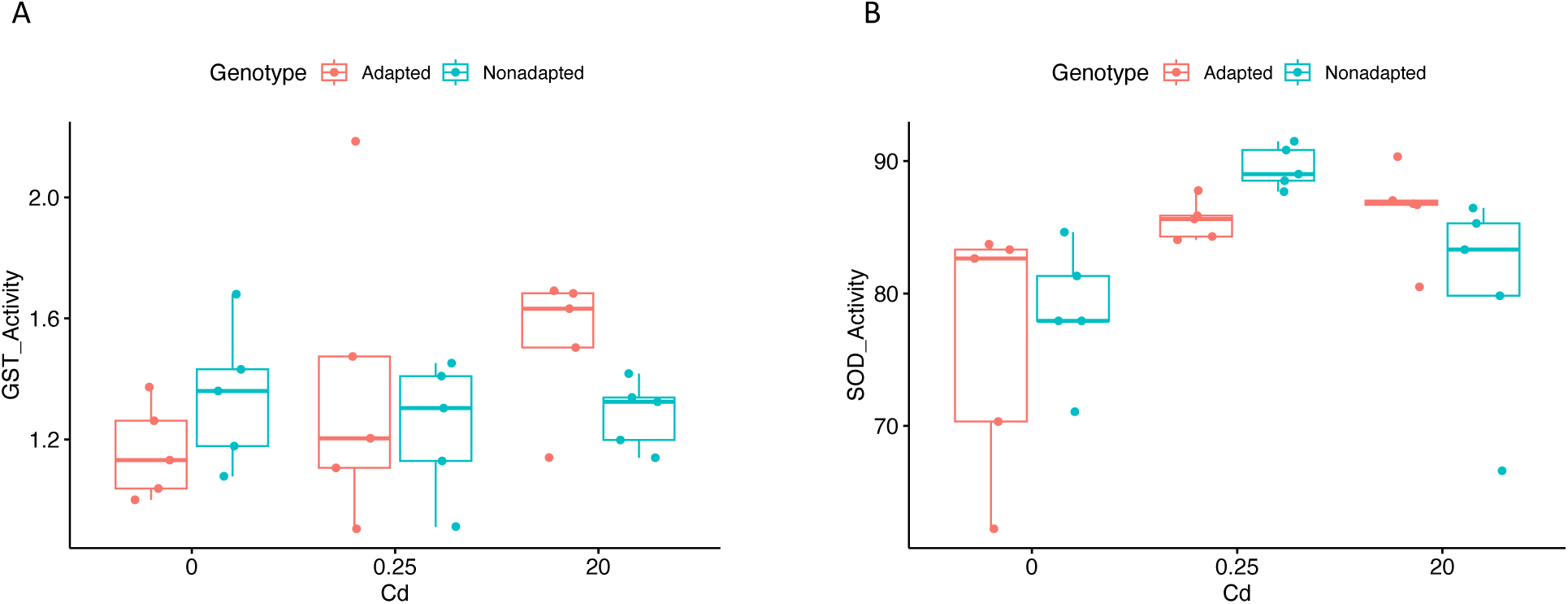
Reproductive rates (r) of adapted and nonadapted MA lines across duration of the experiment.

### Superoxide Dismutase (SOD) Activity

We measured SOD activity in control conditions (0 μg Cd / L) and cadmium exposure (0.25 and 20 μg Cd / L) with both the ADAP and NONA genotypes (Materials and Methods). In ADAP, compared to control conditions, cadmium exposure increased SOD activity at concentrations of 0.25 and 20 μg Cd / L (T-test, P = 0.04 and 0.03 for 0.25 and 20 μg Cd / L, respectively, Fig. 3B). In NONA, relative to control conditions, cadmium significantly increased SOD activity at 0.25 μg Cd / L (T-test, *P* = 8.47 x 10^-4^), but not at 20 μg Cd / L (T-test, *P =* 0.35). Additionally, at 0.25 μg Cd / L GST activity was significantly higher in NONA compared to ADAP (T-test, *P =* 0.002). In 20 μg Cd / L, GST activity was higher in ADAP than NONA, although marginally non-significant (T-test, *P =* 0.08) (Fig. 3B).

### Genome-wide single-nucleotide mutation (SNM) patterns

We observed genome wide SNM rates of 2.06 and 1.74 x 10^-9^ base pair^-1^ generation^-1^ for ADAP control and ADAP cadmium, respectively. These values are similar to the SNM rates reported from other *D. pulex* studies (2.30 x 10^-9^, (Flynn, et al. 2017); 3.35 x 10^-9^ and 4.53 x 10^-9^, (Keith, et al. 2016); 1.57 x 10^-9^ (Keith et al. 2021)). In ADAP, although cadmium exposure did not significantly change the overall SNM rate, cadmium decreased the Ts/Tv ratio (ADAP Control = 1.07, ADAP Cd = 0.93) and increased the G/C → A/T mutation bias (Control = 1.8, Cd = 2.8; Supplemental Tables S3, S5).

In NONA, as we observed in ADAP, cadmium exposure did not significantly change the overall SNM rate (Supplemental Tables S5, S9). However, contrary to our findings in ADAP, cadmium exposure decreased the Ts/Tv ratio and increased G/C → T/A mutation bias in NONA (Supplemental Tables S2, S4).

### Conditional SNM rates

There are six classes of SNMs, with each of these classes originating at either A/T, or G/C positions (e.g., A:T → G:C SNMs originate at A/T positions). The conditional SNM rate of a given SNM class is the per generation rate normalized by the number of genomic positions where the class can arise. In ADAP, cadmium exposure decreased the conditional A:T → G:C mutation rate (T-test, *P*=0.05), and increased the rates of C:G → G:C SNMs, although this difference was marginally non-signficant (T-test, *P=*0.07(Fig. 4A). Additionally, cadmium exposure decreased the rate of G:C → T:A, but only in multinucleotide mutation clusters (“MNMs”, or >2 SNMs within 50 bp) (Fig. S2).

**Figure 4.**
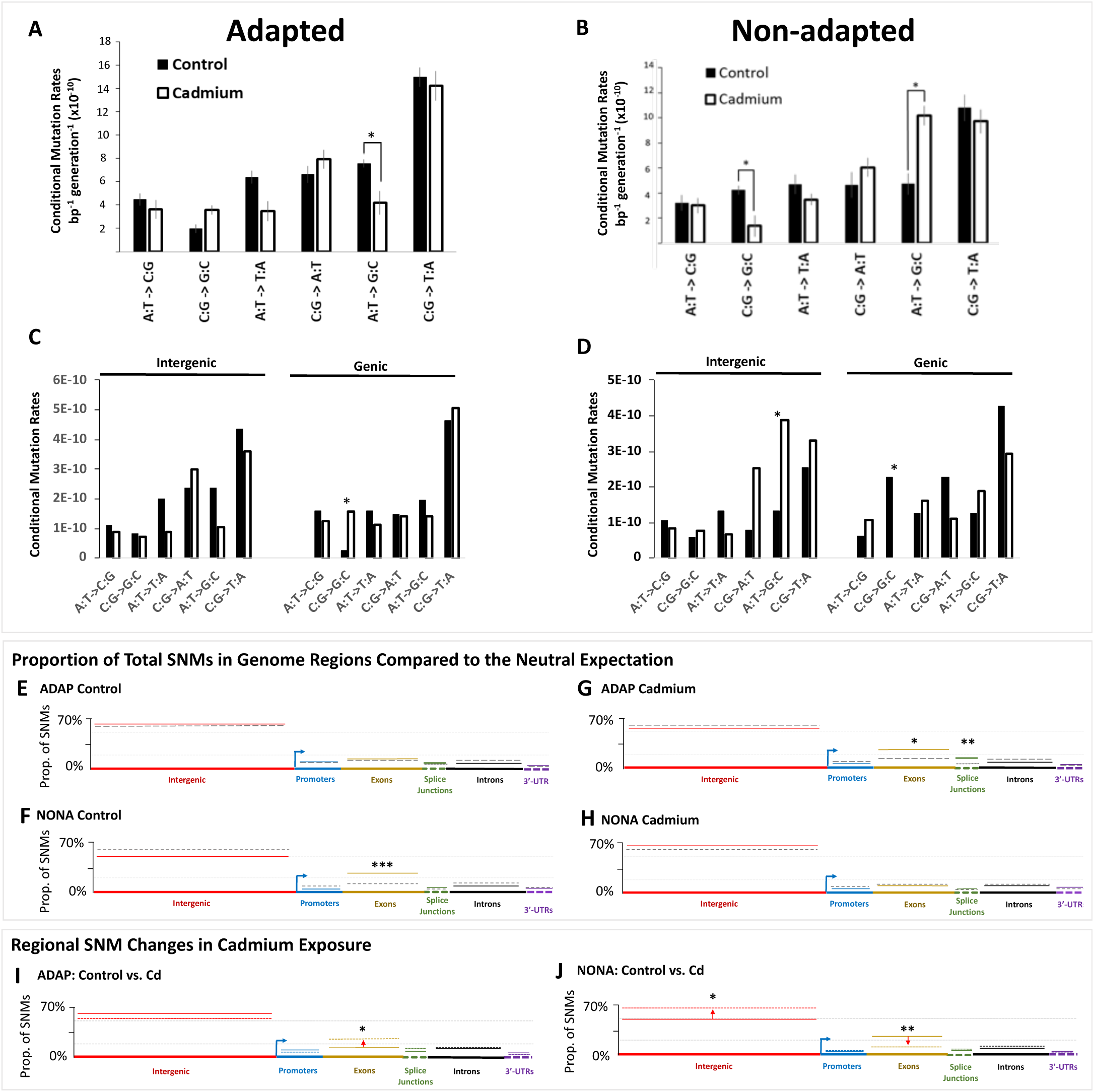
SNM rates in control and cadmium exposure for adapted and non-adapted genotypes. (A, B) Genome-wide conditional SNM rates of control and cadmium exposure. For control (solid black boxes) and cadmium exposure (white boxes), the rates of each of the six classes of base-substitution are plotted. Error bars are included (grey). Significant differences between control and cadmium denoted by asterisks. (C, D) Conditional SNM rates for intergenic and genic regions. The conditional mutation rates are plotted for intergenic (red) and genic (black) regions for the six mutation classes. Solid boxes represent control condition results and white boxes represent cadmium exposure results. Significant differences denoted by asterisks. (E-H) The proportion of total mutations in each (solid lines) compared to the expectation if mutations were randomly distributed across the genome (grey dashed lines). Asterisks denote regions where significant differences were observed. (I, J) Comparison of the proportion of total mutations in each region of control and cadmium exposure for the non-adapted genotype “I” and the adapted genotype “J”. In each region, the proportion of total mutations for control conditions are solid lines, and proportion of total mutations for cadmium exposure are dashed lines. Asterisks denote regions where significant differences were observed.

SNM classes for which the rate was changed by cadmium exposure in ADAP were also changed by cadmium in NONA. However, the direction of cadmium-induced changes in NONA were opposite to ADAP. In NONA, cadmium exposure increased the rate of A:T → G:C, decreased the rate of C:G → G:C, and increased the rate of clustered G:C → T:A MNMs (Keith et al. 2021; Figs. 4B, 4D, Fig. S2).

### Intergenic vs genic mutational patterns

Contrary to ADAP, in NONA, cadmium exposure increased the rate of A:T → G:C in intergenic regions (Fisher’s Exact Test, *p =* 0.04) (Fig. 4D). Further, in ADAP, the elevated, overall rate of C:G → G:C in cadmium exposure was driven by a gene-specific elevation of the C:G → G:C SNM rate (Fisher’s Exact Test, *p =* 0.03) (Fig. 4C), which was not observed in NONA.

### Regional SNM rates

Keith et al. (2021) reported that cadmium exposure changed the regional distribution of SNMs (i.e., intergenic, promoters, exons, introns, splice-site junctions, and 3’UTRs). We therefore investigated cadmium-induced changes to SNMs within these specific genome regions by comparing the proportion of total SNMs in these regions to the expected proportion assuming random distribution across the genome. In ADAP Control there was no difference from the random expectation in any region (Fig. 4E, Table S14). However, in cadmium exposure the proportion of total SNMs in exons (exact binomial test, *P* = 0.03) and intron splice-site junctions (exact binomial test, *P* = 2.20 x 10^-3^) were higher than the random expectation (Fig. 4G, Table S14). Again, we observed opposite trends in NONA compared to ADAP. In NONA control conditions, there are more mutations than expected in exons (exact binomial test, *P* = 3.0 x 10^-4^), and in cadmium exposure no region differed from the random expectation (Figs. 4F, 4I, Table S14).

When the regional proportions of total SNMs of ADAP Control were compared to ADAP Cd (instead of the random expectation), cadmium exposure significantly elevated SNMs in exons (Fisher’s Exact Test, *P* = 0.04, Fig. 4I). Again, the results are opposite in NONA. When we compare the proportions of mutations in NONA Control to NONA Cd, SNMs are significantly decreased in exons (Fisher’s Exact Test, *P* = 4.4 x 10^-3^, Fig. 4J), but additionally, SNMs are also significantly elevated in intergenic regions (Fisher’s Exact Test, *P* = 0.03, Fig. 4J, Table S15).

### Mutation contexts

We next analyzed the mutation contexts of SNMs, the nucleotide where a SNM originated and the directly flanking 5’ and 3’ nucleotides, which can be used to link specific polymerases that induce specific mutation contexts (Supek and Lehner 2017; Keith, et al. 2021). In ADAP Cd all contexts observed less than the neutral expectation were associated with the error-prone, translesion polymerase, Pol η (Supek and Lehner 2017) (χ^2^, *P* = 0.02 (ATT,), *P* = 0.02 (AAT); *P* = 0.04 (TTT); Fig. 5A, 5B; Supplemental Table S11, S13). All significantly elevated contexts are associated with 5-methylcytosine (Zhu, et al. 2014) (5-mC) (χ^2^, *P* = 1.00 x 10^-3^ (CCG); *P* = 0.02 (AGC); *P* = 0.02 (GCT); *P* = 0.03 (GGC); Figs. 5A, 5B; Supplemental Table S16). In contrast to ADAP Cd, in NONA Cd, a Pol η-linked mutation context was significantly elevated compared to the random expectation (χ^2^, *P* = 0.02 (GTT); Figs, 5E, 5F; Supplemental Table S12).

**Figure 5.**
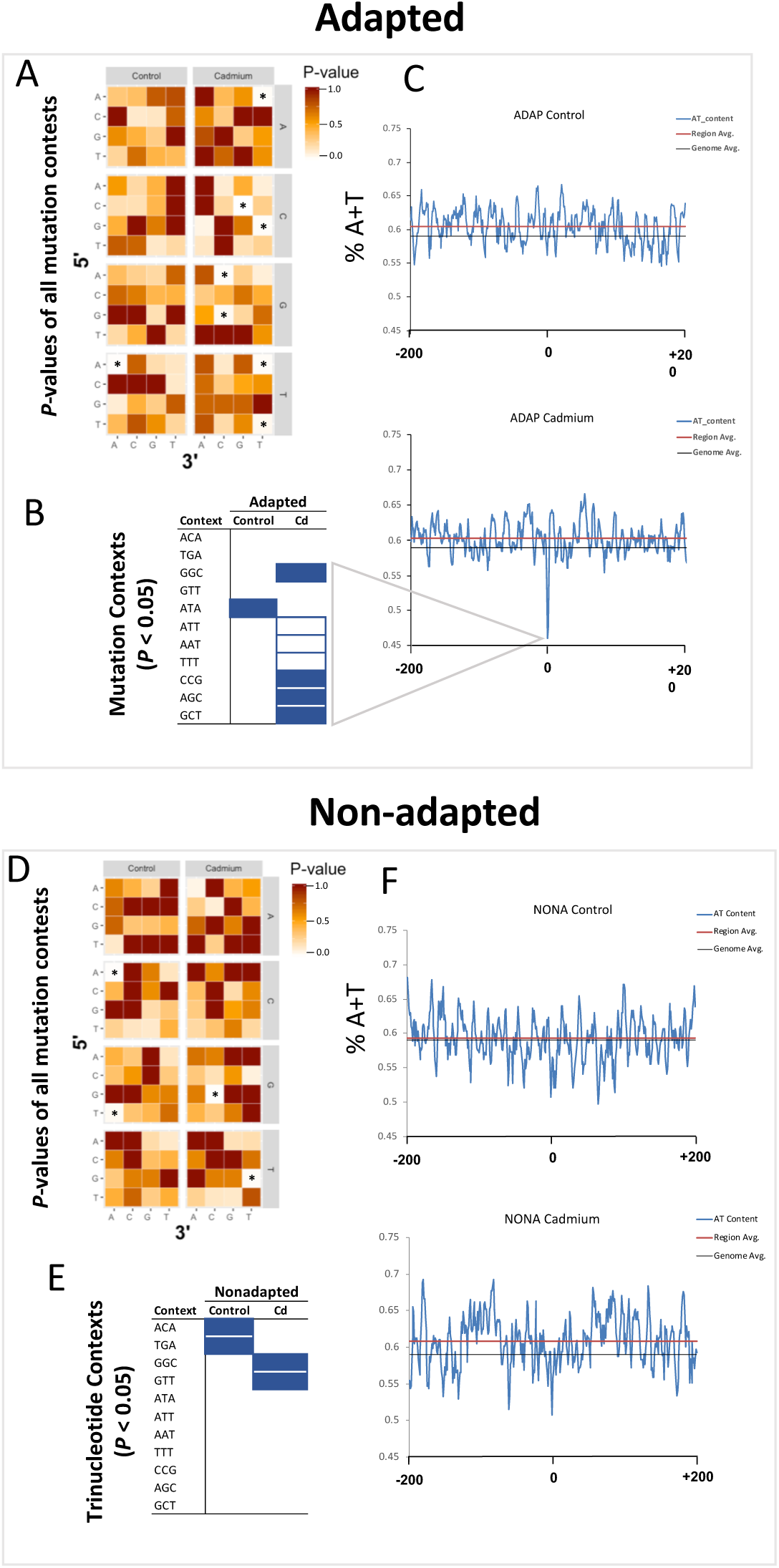
Genomic context of SNMs in non-adapted and adapted genotypes. (A, E). Heat map of *P-*values for all possible SNM contexts of Adapted (ADAP) (B) and nonadapted (NONA) (F) genotypes. The Y-axis is the nucleotide directly 5’ to the SNM. The X-axis is the directly adjacent 3’ nucleotide to the site of SNM. The Z-axis is nucleotide that was changed by SNM. Asterisks denote mutation contexts where *P* < 0.05. B, F. Mutation contexts where *P* < 0.05 for Adapted (B) and Non-adapted (F) genotypes. Solid boxes are contexts where more SNMs were observed compared to the random expectation. White boxes are contexts where less SNMs were observed compared to the random expectation. C, D, G, H. Average A+T nucleotide percentage in the regions around SNMs. For each genotype, and experimental MA condition, SNMs and the −200 and +200 nucleotides were “stacked” resulting in a 401-nucleotide alignment of each SNM and their surrounding region. A four nucleotide “sliding window” moved from −200 to +200 in 1 nucleotide increments on this alignment. For each 4-nucleotide window (from −200 to +200), the average A+T% across of all aligned SNM regions are plotted, resulting in a visualization of consensus A+T% at each window of SNM regions.

### 5-hydroxymethylcytosine Analysis

We previously reported that cadmium reduces 5-hydroxymethylcytosine (5-hmC) levels in NONA. We therefore investigated 5-hmC levels in cadmium in ADAP. In ADAP, compared to control, cadmium does not significantly reduce 5-hmC levels at 0.5 μg Cd / L (T-test, *P =* 0.29) or 20 μg Cd / L (T-test, *P* = 0.18) (Fig. 6A). Contrary to ADAP, in NONA, cadmium did significantly reduce 5-hmC levels compared to control at 0.5 μg Cd / L (T-test, *P =* 0.01) and 20 μg Cd / L (T-test, *P* = 0.02) (Fig. 6B).

**Figure 6.**
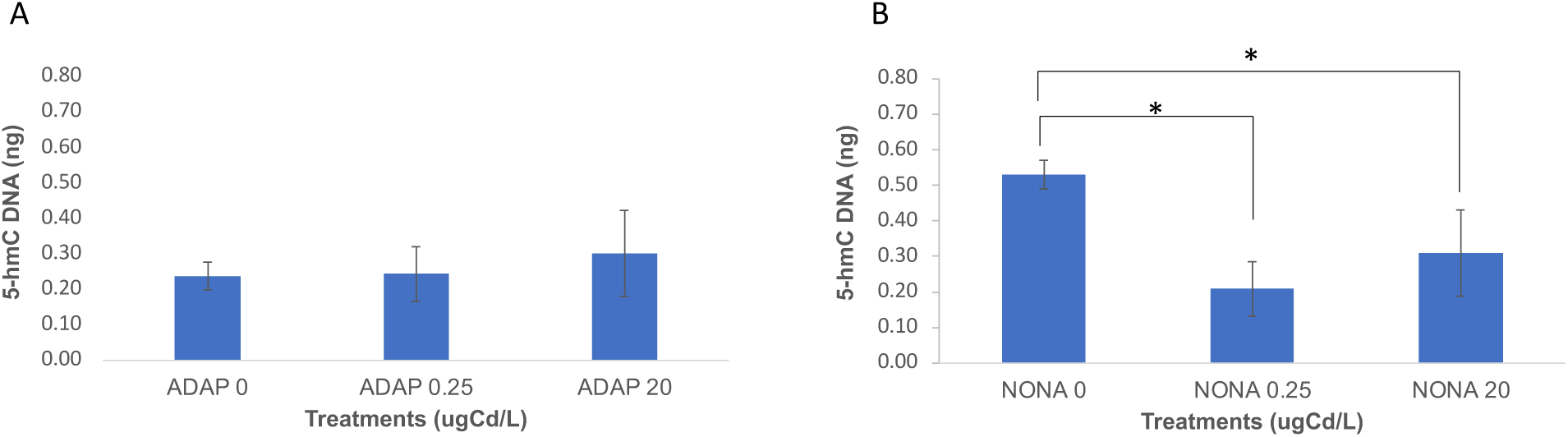
(A) 5-hydroxymethylcytosine levels in ADAP at concentrations of 0, 0.25, and 20 μg Cd /L. The mean and corresponding standard error of the mean are plotted. (B) 5-hydroxymethylcytosine levels in ADAP at concentrations of 0, 0.25, and 20 μg Cd /L. The mean and corresponding standard error of the mean are plotted.

### de novo CNV Analysis

We identified de novo CNVs > 3,000 bps (i.e., spontaneous CNV mutations that occurred during the MA experiment) according to our previously published methods outlined in Keith et al. (2015). In ADAP Control, 65 de novo CNVs were identified and 59 CNVs were observed in ADAP Cd. CNVs ranged in length from 3 kb to 973 kb in ADAP Control, and 3 kb to 801 kb in ADAP Cd. Average de novo CNV length did not differ between ADAP Control and ADAP Cd (Mann-Whitney U, *P* = 0.80) (Table S18).

Notably, a large proportion of ADAP Control de novo CNVs are observed in a single MA subline (ADAP Control 10, which is 1 out of 12 analyzed sublines in the ADAP Control experiment). This subline contains 62% (42 of 68) of the observed de novo CNVs found in the ADAP Control MA experiment (12 MA sublines), and it contains 47% of the base-pairs subjected to de novo CNVs in the ADAP Control MA experiment (Table S18). Over half of the de novo CNV base-pairs in ADAP Control 10 are located on Chromosome 3 (57%) and are scattered across eight scaffolds (Table S18).

The overall number and rates of de novo CNVs observed in NONA Control and NONA Cd are over an order of magnitude lower than what is observed in ADAP Control and ADAP Cd (Figs. S5 and S6). In NONA Control, only 9 de novo CNVs were observed ranging from 3,500 - 212,500 bps. In NONA Cd, only 4 de novo CNVs were observed ranging from 4,000 - 13,500 bps.

### Metallothionein-1 Copy-number Analysis

Metallothionein (*MT1*) is a protein that protects organisms against cadmium toxicity by irreversibly binding and sequestering cadmium (Shaw et al. 2006). We therefore investigated *MT1* copy-number and gene expression levels in cadmium-adapted and non-adapted populations, including the adapted and non-adapted genotypes, with quantitative PCR (qPCR).

*MT1* gene copy-number is significantly higher in adapted genotypes than nonadapted genotypes (*P* = 0.001, Fig. 5C, Fig. 7C, Table S19). Basal (control) *MT1* gene expression is also significantly elevated in adapted genotypes compared to nonadapted genotypes (T-test, *P* = 2.83 x 10^-5^) (Table S19). Cadmium exposure increased *MT1* expression in both adapted (T-test, *P* = 1.26 x 10^-8^) and nonadapted (T-test, *P* = 3.09 x 10^-10^) populations. However, *MT1* expression was induced to higher levels in adapted genotypes, although marginally non-significant (T-test, *P* = 0.06). Further, we observe a strong correlation between *MT1* expression and gene copy-number in adapted genotypes (Figure 7D, *R =* 0.8, p = 0.016). However, no correlation between *MT1* expression and gene copy-number is observed in nonadapted genotypes (Figure 7D, *R* = 0.0023, *p =* 1).

**Figure 7.**
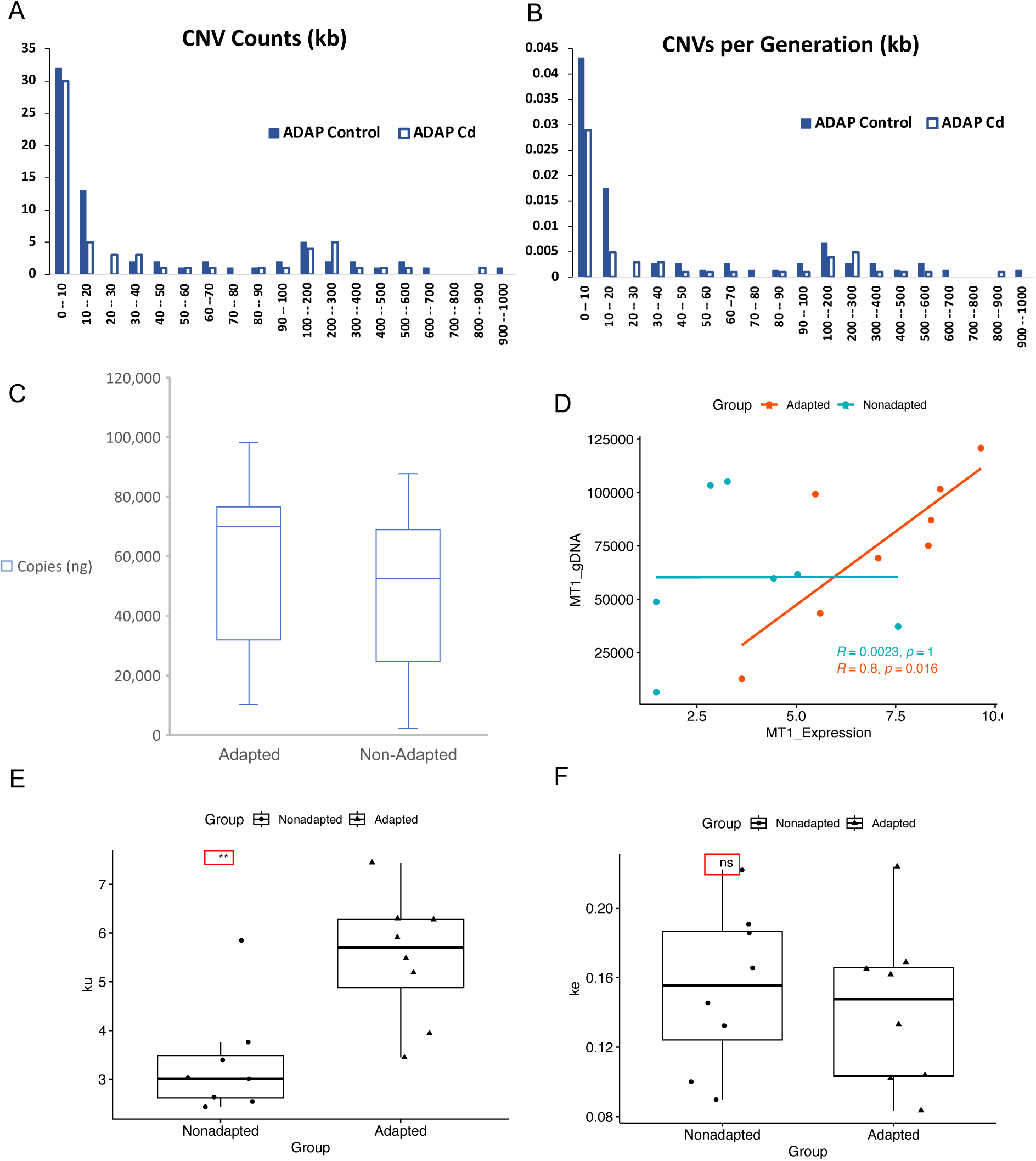
(A) Size distribution of de novo CNVs in ADAP Control and ADAP Cd. (B) CNV rate (CNVs per generation) in ADAP Control and ADAP Cd. (C) qPCR measurements of metallothionein (MT) gene copy number in adapted and nonadapted populations. Y-axis is the number of qPCR amplicons (ng) (Methods). (D) Correlation of MT1 gene copy number and RNA expression for adapted and nonadapted *D. pulex* lake populations. (E) Cadmium elimination results for adapted and nonadapted populations (k_e_). (F) Cadmium uptake (k_u_) results for Adapted and Nonadapted populations.

### Cadmium Uptake and Elimination Analysis

If the higher copy-numbers of *MT* and the corresponding higher gene expression level is an explanation for protection against cadmium-induced mutagenesis in adapted genotypes, then the tissues of adapted populations will have higher levels of MT-bound cadmium compared to nonadapted populations. Consistent with this expectation, the rate of cadmium uptake was 65% higher in adapted populations compared to nonadapted populations (adapted, k_u_* = 5.5 ± 1.3 µg Cd g^-1^ dry wt day^-1^, n=8, non-adapted, k_u_* = 3.3 ± 1.3 µg Cd g^-1^ dry wt day^-1^, n=8, mixed liner model, P<0.01), while the elimination rate for adapted populations (k_e_ = 0.14 ± 0.05 day^-1^, n=8) and non-adapted populations (k_e_ = 0.15 ± 0.05 day^-1^, n=8) did not differ. The increased uptake in the absence of compensatory decreases in cadmium elimination resulted in increased cadmium burden in adapted populations compared to nonadapted populations (adapted 13.1 ± 3.1 µg Cd g^-1^, n=8, nonadapted 7.3 ± 2.1 µg Cd g^-1^, n=11, mixed liner model, P<0.01). These data are consistent with superior cadmium sequestration in the cadmium adapted isolates (Shaw, et al. 2007; Asselman, et al. 2013).

## Discussion

After extensive lake sampling of *D. pulex* populations from a region devastated by industrial mining and a nearby region without a history of mining, we report that many *D. pulex* lake populations from Sudbury are adapted to cadmium. We used MA experiments to compare the effect of cadmium on germline mutation between adapted and non-adapted genotypes and found that cadmium exposure altered the SNM rate in the same genome regions, but in opposite directions. Cadmium also changed the same rates of specific SNM classes (A:T → G:C and C:G → G:C), albeit in opposite directions for adapted and non-adapted genotypes. The reversal of germline mutational trends in the adapted MA experiment supports the hypothesis that adaptation to cadmium protects the germline from toxicant-induced mutagenesis.

Our findings reveal that in the NONA (nonadapted) MA experiment, fitness in cadmium exposure lifetables was lower than control condition lifetables, while the opposite was observed in the ADAP MA experiment – i.e., in the cadmium-adapted, genotype lifetable fitness over the course of the experiment was higher in cadmium than in control conditions. Adaptation occurs when natural selection, retains tolerant genotypes (Hochmuth, et al. 2015). Adapted phenotypes are therefore expected to have higher fitness when subjected to the condition(s) in which they are adapted.

Multiple studies have investigated the fitness of adapted animal phenotypes in the presence and absence of a chemical stressor or environmental condition (Hochmuth, et al. 2015; Flynn, et al. 2019; Shaw, et al. 2019; Pham, et al. 2021) – showing that the adapted phenotype has higher fitness in the stressor or condition to which they are adapted. However, our study and results are different, as this study directly measures fitness in adapted and nonadapted genotypes under strict genetic bottlenecks, which therefore measures relative fitness increases and declines that are the direct result of de novo genome-wide mutations in the experiment. Our fitness assay in the adapted genotypes (ADAP) clearly shows there is less of a genome-wide mutational impact on fitness in the presence of the chemical to which it is adapted, and that a secondary consequence of adaptation to a chemical stressor or environmental condition is that the mutations accrued have less of a fitness impact.

Results from MA experiments with the nonadapted genotype (NONA) suggest that cadmium induced oxidative stress and substitution with zinc domains within proteins contribute to the observed cadmium-induced mutagenesis (Keith, et al. 2021). In NONA, we previously suggested that the cadmium-induced increase of the A:T → G:C SNM rate by interfering with DNA Polymerase η (Keith, et al. 2021). Pol η contains a zinc-finger domain whose structure is essential for initiation of translesion synthesis (Bienko, et al. 2010). Cadmium readily inhibits zinc-finger domains because of cadmium’s higher affinity compared to zinc for the thiol groups of cysteine (Li and Manning 1955; Hartwig 2001). Our results therefore suggested that in NONA, cadmium inhibits Pol η function and increases the A:T → G:C SNM rate.

In ADAP, we observed a decrease in the A:T → G:C rate in cadmium exposure (Fig. 4A), which would be expected if cadmium adaptation protects the germline from inhibition of Pol η. Giving further evidence for protection in ADAP against Pol η-induced SNMs are the mutational context findings. In ADAP Cd, all contexts observed less than the neutral expectation are associated with Pol η (Figs. 5A, 5B).

In NONA we suggested that the decreased C:G → G:C rate in cadmium exposure was a result of cadmium inhibiting the two, zinc-finger domains of TET proteins (Hu, et al. 2013), which convert 5-methylcytosine (5-mC) to 5-hydroxymethylcytosine (5-hmC) (Ito, et al. 2011). Because 5-hmC positions were recently determined to have elevated rates of C:G → G:C SNMs (Supek and Lehner 2017), we reasoned that cadmium inhibits TET zinc-finger domains, which reduced 5-mC → 5-hmC conversion, and lowered the rate of C:G → G:C SNMs.

Contrary to NONA, in ADAP we observed an increased C:G → G:C SNM rate in cadmium exposure relative to control - suggesting cadmium adaptation protects TET from cadmium inhibition, allowing for 5-mC → 5-hmC conversion, and ultimately, mutation. Here, we further show that while in NONA, Cd exposure significantly reduces 5-hmC, 5-hmC levels are maintained in ADAP under Cd exposure (Fig. 6). Although investigations into the function of 5-hmC is a relatively new field of study, our results show that the maintenance of 5-hmC in ADAP is also maintaining the function of 5-hmC positions, but also results in elevated numbers of C:G → G:C mutations in genic regions.

Due to the extreme low de novo CNV rate in NONA we are not able to make conclusions about cadmium’s effect on CNV occurrence (Figs. S4, S5). In NONA, we only observed 9 de novo CNVs and 4 de novo CNVs in Control and Cd conditions, respectively. However, we do observe a marked decrease to CNV rate in ADAP Cd relative to Control, suggesting lowered CNV rates under cadmium exposure in this adapted genotype (Figs. S4, S5). Notably, from this study and a previous *Daphnia pulex* mutation-accumulation study (Keith et al. 2016), the de no novo CNV rate differs between genotypes by over 23-fold (Fig. S5), ranging from 0.006 (NONA Cd) - 0.140 de novo CNVs per generation (LIN genotype, Fig S5; Keith et al. 2016).

Keith et al. (2016) showed that in the LIN genotype the CNV rate is highest in regions of the genome that are highly heterozygous due to introgression of two chromosomes from a genetically divergent sister species (*D. pulicaria*) (Keith, et al. 2016). We therefore measured genome-wide levels of heterozygosity (*π*_t_) for NONA, ADAP from this study, as well as genotypes LIN and TCO which we previously measured de novo CNV rates for (Keith et al. 2016), and compared these values to de novo CNV rates. Although there are only four genotypes, the two genotypes with the highest *π*_t_ (LIN, *π*_t_ = 0.023, and ADAP *π*_t_ = 0.026) also have the highest de novo CNV rates, while NONA (*π*_t_ = 0.015) and TCO (*π*_t_ = 0.002) have lower de novo CNV rates (Figure S5).

We identified elevated pre-existing copy-numbers and gene expression levels of *MT1* in cadmium-adapted populations compared to nonadapted populations. First identified as a cadmium-binding protein in 1957 (Margoshes and Vallee 1957), MT is a small protein (∼6,500 Daltons) with many unusual characteristics, including lack of a secondary structure or aromatic amino acids (Vallee 1991; Palmiter 1998). An analysis of the *MT* amino acid sequence across Crustacea showed that *MT* is non-conserved except for a high percentage of cysteine amino acids (30-33% of total *MT* amino acids (Shaw, et al. 2007). The conserved cysteine amino acids of *MT* coordinate non-reversible MT-Cd binding. The increased levels of MT in adapted populations (including the MA genotype ADAP) would therefore be expected to reduce cadmium reactivity within cells, decrease oxidative stress, and prevent zinc inhibition by cadmium in zinc-containing protein domains.

Increasing MT copy number and expression level is known to be one path to Cd adaptation. *MT* was duplicated in evolved, cadmium-adapted *Drosophila melanogaster* strains, which correspondingly elevated *MT* expression by two-fold (Otto, et al. 1986). Another study that analyzed globally distributed *Drosophila* strains showed that genotypes with multiple *MT* copies were more tolerant to cadmium compared to those without *MT* duplications. *MT* gene expression was increased in the strains with *MT* duplications (Maroni, et al. 1987), as observed by (Otto, et al. 1986). Our identified *D. pulex* genotypes from adapted populations (including the genotype ADAP) have increased copies of the metallothionein-1 gene relative to nonadapted from Dorset, which results in increased basal levels of its expression and increased levels following induction with Cd in adapted genotypes. These attributes are expected to enhance sequestration of cadmium, which was confirmed through direct measurements of cadmium uptake, elimination, and burden in cadmium-adapted and non-adapted *Daphnia*.

Our finding that in the adapted genotype DNA repair mechanisms function more efficiently in cadmium exposure has important implications for determining environmental remediation strategies. In a polluted environment where populations have adapted, if the rate of remediated occurs swiftly, it can remove adapted populations from their optimal environments before they can re-optimize, if possible, to pristine conditions. The reversal of mutational trends we report here therefore argues for phenotypic considerations when enacting remediation strategies.

## Materials and Methods

### Hybrid Vigor Analysis

We utilized 17 *D. pulex* × *D. pulicaria* F_1_ hybrids produced in the lab for a previous study (Heier and Dudycha 2009), 9 North American *D. pulicaria* strains and 5 North American *D. pulex* strains, 10 of which were sires or dams, respectively, of the F_1_ hybrids. *D. pulicaria* and *D. pulex* strains, as well as the F_1_ hybrids, were obtained from the *Daphnia* collection of the Michael Lynch Lab, Biodesign Institute Center for Mechanisms of Evolution, Arizona State University, Tempe. *Daphnia* strains were maintained in COMBO medium (Kilham et al. 1998) at 20° C in the laboratory with a 16:8 hour light:dark cycle. Strains were fed *Ankistrodesmus falcatus* every 2 days during bulking and culturing to obtain experimental neonates, with at least 3 generations before experimentation. Then 21-day cadmium exposure experiments were performed on even-aged neonates individually in 0, 2.5, or 5 µg/L Cd conditions with 5× replication for each condition, following chronic exposure protocols similar to Asselman et al. (2013). Strains were fed *A. falcatus* daily and transferred to new media for observation and counting of neonates, which were then removed. Total reproduction was calculated by summing the total offspring of an individual during its lifespan.

To analyze the effects of hybridization and cadmium stress on total reproduction in F_1_ hybrids and their parental strains, we fitted a linear mixed-effects model (LMM) using the lme4 (v1.1_33) R package (Bates et al. 2015). The model included the fixed effects of genotype group (*D. pulicaria, D. pulex,* F_1_ hybrid) and cadmium concentration (0, 2.5, 5 µg/L), and their interaction. The random effects structure included strain nested with genotype group to account for the hierarchical design of the experiment, where there were multiple strains with replication within each genotype group. This model explicitly partitions variance among strains and among replicates within strains, preventing pseudoreplication. Significance of fixed effects was assessed using Type III Analysis of Variance using the lmerTest (v3.1_3) R package (Kuznetsova et al. 2017). Post-hoc analyses were conducted using the emmeans (v1.8.6) R package (Searle et al. 1980, Lenth 2023). To test for hybrid vigor, we computed a contrast comparing the F_1_ hybrid mean to the mid-parent mean within the control condition. To evaluate fitness at each cadmium concentration, we examined the simple effects of the significant interaction by performing Tukey-adjusted pairwise comparisons between genotype groups within each cadmium concentration. Finally, to test for enhanced cadmium tolerance in hybrids across all cadmium concentrations, we tested for differences in the linear slopes of reproductive decline between the F_1_ hybrids and each parental species. Plots were generated using ggplot2 (Wickham 2016) and all statistics computed in R v4.1.3 (R Core Team 2022).

### Reproductive rate (r) analysis

Chronic life table tests following ASTM methods (ASTM 1990) and detailed in Shaw et al. (2019) were performed at generations 10, 25, and 40, respectively. Reproductive rate (r) was determined by summing the daily reproductive daily output (as described in (Chen and Folt 1996)) for each of five clonal replicates of 90 different MA line replicates in 0 μg Cd/L (methods detailed in (Shaw, et al. 2007)). The test length was modified from the traditional 21 days to 30 days to adjust for the life-history dynamics of the ADAP and NONA genotypes, which ensured life-history data were collected from three separate broods.

### GST Activity Assay

GST activity was measured with the Glutathione S-Transferase (GST) Assay Kit from Sigma (Cat. No. CS0410). In this method, 1-Chloro-2,4,dinitrobenzene (CDNB) is utilized, as it provides a broad range of GST isozyme measurement. CDNB conjugation with glutathione results in an increased absorbance at 340 nm. The increase in absorption at 340 nm is proportional to GST activity. We followed *Daphnia* methods outlined by (Oexle, et al. 2016), with the exception that we normalized final GST activity to volume of supernatant that was recovered from large, glass tubes that were used to grind daphnia, which provided a precise measurement of the amount of input supernatant into the final reaction. 195 µL of substrate solution and 5 µL sample, GST, or control was added per well. Absorbance was measured every minute for 7 minutes. For the NONA and ADAP genotypes, we used 5 biological replicates with two technical replicates for Cd exposures of 0 μg Cd/L, 0.25 μg Cd/L, and 20 μg Cd/L.

### SOD Activity Assay

SOD activity was measured with the Superoxide Dismutase (SOD) Activity Assay Kit from Sigma-Aldrich (Cat no. CS0009) per manufacturer’s recommendations. SODs catalyze the conversion of superoxides to O_2_ and H_2_0_2_. This method uses xanthine oxidase to produce superoxide anions. The levels of superoxide anions then interact with WST dye forming formazan dye, which can be measured at 450 nm. Reduction in measurement of formazan dye at 450 nm is the result of increased SOD(s) activity in the sample. A standard curve, blanks, and controls were prepared according to manufacturer’s recommendations. 20 µL of sample was added to each well followed by 20 µL Dilution Buffer and 160 µL of WST working solution. 20 µl xanthine oxidase was used to initiate the reaction. For the NONA and ADAP genotypes, we used 5 biological replicates with two technical replicates for Cd exposures of 0 μg Cd/L, 0.25 μg Cd/L, and 20 μg Cd/L.

### MA line maintenance

Methods for MA line maintenance were described in (Keith, et al. 2021). Briefly, for each subline, after clonal reproduction each generation one offspring was randomly selected within 24 h from reproduction to serve as the mother for the next generation. We additionally randomly selected two offspring (for each subline) which served as backups. Exposure water (control and Cd) was changed each generation and Cd was measured throughout the experiment to ensure concentrations were as expected (Dartmouth Trace Element Analysis Core). Sublines were maintained in laboratory culture as described by (Shaw, et al. 2007). *Ankistrodesmus falcatus* (75,000 cells/mL) was fed to the sublines, daily. Sublines were maintained under standard light:dark laboratory conditions (20°C; 12 h light: 12 h dark). The control condition was maintained in 50-mL beakers of modified COMBO media, lacking nitrogen and phosphorus (Kilham, et al. 1998). Cd exposure sublines were maintained in modified COMBO media at a concentration of 0.25 μg Cd/L (Cadmium Chloride, ACS/analytical grade; Fischer Scientific) (Shaw, et al. 2007). Primary stocks of Cd were made by dissolving CdCl_2_ (analytical grade, Fischer Scientific) in ultrapure water. Test Cd concentrations were verified every year at the Dartmouth Trace Element Analysis Core with a magnetic sector inductively coupled plasma/mass spectrometer (ELEMENT; ThermoElectron) fitted with a standard liquid sample introduction system (microconcentric nebulizer (MCN–2; CETAC) and cooled Scott-type spray chamber) (Shaw, et al. 2019).

### Read Processing and Mapping

Next-generation sequencing read processing was performed as previously described by (Keith, et al. 2021). Briefly, paired-end read trimming was performed with Trimmomatic (Trimmomatic v.0.36, USADEL Lab) (Bolger, et al. 2014) in Paired-end mode with parameters ILLUMINACLIP:2:30:10 LEADING:28 TRAILING:28 MINLEN:50. Mapping was performed with BWA-MEM (BWA-MEM v0.7.17, Heng Li Lab) (Li and Durbin 2009) to the *Daphnia pulex* reference assembly V1.0 (GCA_0001878751.1) using paired-end mode and default parameters. We generated four SAMtools (Samtools v1.9, Wellcome Sanger Institute) (Li, et al. 2009) mpileup files: NONA control, NONA Cd, ADAP control, and ADAP CD. These mpileups reported only high-confidence genotype calls (parameters –Abx –min-BQ 30 –min-MQ 30).

### Mutation Identification

We previously published and provided an openly available python program for calling mutations in deep-coverage, whole-genome-sequenced MA experiments (Mutation_Caller.py,(Keith, et al. 2021)). Mutation_Caller.py was used here to call mutations in ADAP control and Cd. The following criteria are used by Mutation_Caller.py to identify mutations. a) For a site to be analyzed a minimum proportion of mapped reads was 0.9 for homozygous sites, and between 0.3 for and 0.7 for heterozygous sites. If a given site met either of these criteria across all sublines in either control or cadmium conditions, and if all sublines had the same genotype, then this shared genotype was denoted as the “consensus” genotype. Additionally, if all but one subline shared a given genotype, then this unique genotype was considered a potential mutation, or “putation”. b) A minimum sequencing depth of coverage of 12× and a maximum depth of coverage over 45×. c) 20 bp around indels (i.e., indels between our genotype and the *Daphnia pulex* reference genome) were excluded. d) A minimum of two reads were mapped in both orientations supporting each genotype call to eliminate false genotype calls resulting from PCR artifacts. e) Reads that mapped to multiple loci were removed to eliminate repetitive regions.

### Mutation Rate Calculation

Mutation rates were calculated according to the methods outlined by (Keith, et al. 2016; Keith, et al. 2021). For both NONA and ADAP, for each subline in control and Cd exposure, we independently calculated the genome-wide SNM rate with μ_*bc*_=mi/(2niT) (Lynch, et al. 2008; Sung, Tucker, et al. 2012b), where μ*_bc_* is the genome-wide SNM rate, *m* is the total number of SNMs, 2*n*_*i*_ is the total number of analyzed diploid sites, and *T* is the total number of generations the given subline was propagated. The standard error (SE) for each subline was calculated with 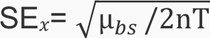, where μ_*bc*_ is the SNM mutation rate for a given subline, *T* is total number of generations for the subline, and *n* is total number of sites analyzed for the subline. For both NONA and ADAP, independently for control and Cd exposure, the overall genome-wide SNM rate was calculated as the average of the genome-wide SNM rates of all sublines. The overall SE was calculated independently for NONA and Cadmium, control and cadmium exposures with 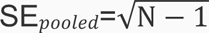, where *s* is the standard deviation of the SNM rate for all sublines, and *N* is the total number of sublines that were analyzed. For NONA and ADAP control and Cd exposures, independently, for each subline, the conditional mutation rates for each of the six classes of SNMs were calculated with μ_*cond*_ = m_i_/(2n_i_T), but with *n*_*b*,*i*_ instead of *n*_*i*_, and *m*_*b*−>*d*,*i*_ instead of *m*_*i*_, where *n*_*b*,*i*_ is the number of ancestral sites of nucleotide type *b* (*b* = A, T, G, or C) in an MA line *i*, and *m*_*b*−>*d*,*i*_ is the number of SNMs from nucleotide type *b* to any of the three possible base pairs to which it can mutate. The overall conditional mutation rate for each of the six SNM classes was calculated as the average across all sublines of each SNM class. This was done for NONA and ADAP, control and Cd exposures, independently. For each SNM class, the SE was calculated according to 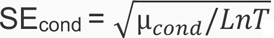, where *u*_*cond*_ is the rate for a specific class of mutation, *L* is the number of lines analyzed in either control and Cd exposure*, T* is the average number of generations, and *n* is the number of analyzed sites. This was performed for NONA and ADAP, control and Cd exposures, independently.

### Genomic DNA MT1 qPCR Methods

High molecular weight genomic DNA was isolated from *Daphnia pulex* clones using the CTAB method and quantified via Nanodrop spectrophotometer (Nanodrop Technologies) and Quant-It PicoGreen (Invitrogen). 4 ng/μl working stocks of genomic DNA template were created and stored at 4°C. 25 μl qPCR reactions were setup using PerfeCTa SybrGreen Mastermix, low Rox (Quanta Biosciences), 20ng of genomic DNA template, and 0.3μM final primer concentrations. Metallothionein I gene was assayed using the following primers: MT1_F 5’ CTTGTCTAACCGATACGTCC 3’ and MT1_R 5’ GTTGAAATGTTGCGAGGG 3’. All primers were ordered from IDT with standard desalting. Thermal cycling was done using a Stratagene MX3000P (Agilent) using the following parameters: 95°C for 7 minutes, followed by 40 cycles of 95°C for 40 seconds, 61°C for 30 seconds, 72°C for 35 seconds, followed by standard dissociation curve.

### MT1 RT-qPCR Methods

RT-qPCR was performed according to the methods outlined by (Asselman, et al. 2012). Briefly, total RNA was extracted with the RNeasy and Qiashredder (Qiagen, Venlo, Netherlands) according to manufacturer’s protocol, followed by DNase treatment (Qiagen, Venlo, Netherlands). First strand-specific cDNA was synthesized using 1 μg RNA with MessageAmpTM II mRNA amplification kit according to the manufacturer’s protocol. RT-qPCR was performed on the Mx3000P Stratagene qPCR system with forward primer 5’-TGGTGGTGAATGTAAATGCAGCGG-3’, and reverse primer 5’-TGTAGTCGTTTACTTGCAGCAGGC-3’. The standard curve consisted of dilutions from a single cDNA sample. Each sample consisted of three biological replicates, with two technical replicates per biological replicate. SYBR Green Super Mix (Quanta Biosciences, Gaithersburg, Md), which contained all PCR components, was used with the addition of the aforementioned primers. 1 μL cDNA was added to the qPCR mixture for a volume of 25 μL total reaction. Following 3 minutes at 95°C, amplification consisted of 40 cycles (30 s at 95°C, 30 s at 60°C, 35 s at 72°).

### Cd Assimilation and Elimination Methods

For each of eight non-Cd-adapted and eight Cd-adapted isolates of *Daphnia pulex*, a total of between 34 and 50 individuals of between twelve and fourteen days old were divided equally over two replicate glass beakers per isolate containing 500 mL of COMBO medium (Kilham, et al. 1998) modified without N or P (Shaw, et al. 2007). The medium also contained a 0.5 µg Cd / L pulse (nominal added concentration), which had been spiked into COMBO prior to transfer of the daphnids and which had been allowed to mix and equilibrate with the medium for 1 hour. Daphnids were then held for 4 days in this medium in a climate room held at between 19.5° and 20.5 °C under a 12h-12h dark-light cycle. Daphnids were fed every other day with *Selenastrum capricornutum*. After 4 days, all daphnids were removed from the Cd exposure medium and transferred for 8 hours to 500 mL of COMBO medium per daphnid, holding fresh food but not containing any Cd, in order to allow depuration of Cd in the gut lumen that was not assimilated in tissue (Gillis, et al. 2005). After this period of 8 hours, half the number of daphnids (the ‘elimination’ or E-daphnids) remained in the clean COMBO medium for another 3 days, while the other half (the ‘assimilation’ daphnids or A-daphnids) were briefly rinsed with 1 mmol/L of Na_2_EDTA to remove potentially remaining carapace-bound Cd. The E-daphnids underwent the same gut depuration and Na_2_EDTA rinse steps as the A-daphnids after a total period of 3 days in the clean COMBO medium. Following the rinsing step, A-daphnids and E-daphnids were dried at 40°C until constant dry weight (24 hours was always sufficient). Dried daphnid pools were then digested in 10mL AAS polyethylene tubes containing 300 µL ultrapure HNO_3_ (Normatom Quality, VWR International, Belgium) with the aid of a microwave. The digests were then diluted 20-fold with ultrapure water and then stored in polyethylene tubes in the dark at 4°C. Prior to analysis on the Graphite Furnace AAS (Thermo Fisher Scientific, Waltham, MA, USA) these samples were diluted an additional 5-fold with ultrapure H2O. A certified plankton reference material [CRM 414 trace elements in plankton, Cd content 0.383 μg/g +/- 0.014] was digested in parallel (9 replicates) to check the accuracy of the analytical method. Reference plankton contained 0.290 ± 0.013 µg/g dry wt (n=12). In addition, procedural blanks were prepared, i.e. acid digests of HNO_3_ not containing any digested daphnid tissue. Procedural blanks contained 0.437 ± 0.303 µg/g dry wt (n=9) and a method detection limit (MDL, as 3 times the standard deviation) of 0.9 ng was derived from that. All measured daphnid samples contained considerably more than the mean of the procedural blanks and also more than the MDL, i.e. A-daphnid samples between 2.1 and 18.3 ng Cd and E-daphnid samples between 1.4 and 9.6 ng Cd.

Internal Cd contents and dry weights of the A- and the E-daphnids were used to estimate uptake and elimination rates of Cd, assuming a one-compartment biokinetic model with first order uptake and elimination kinetics:

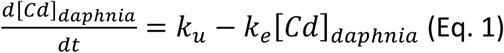

where [Cd]_daphnia_ is the whole-body content of Cd (µg Cd individual^-1^), k_u_ is the first-order Cd uptake rate constant (µg Cd individual^-1^ day^-1^) and k_e_ is the first-order Cd elimination rate constant (day^-1^).

Duration the elimination period (no Cd in exposure medium), k_u_ = 0 and thus integration of (Eq. 1) between the start of the elimination period of the experiment (=end of 4d-exposure period; measured Cd in A-daphnids, [Cd]_daphnia,A_) and the end of the 3-day elimination period (t_elimination_=3d) (measured Cd in E-daphnids, [Cd]_daphnia,E_) gives:

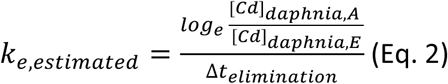

Measurements of A-daphnids and E-daphnids originating from the same exposure beaker were used to calculate one of two duplicate estimates of k_e_ (i.e. one per beaker). If k_e,estimated_ is known, it is possible to account for elimination of assimilated Cd that is already occurring during the exposure period. Furthermore, prior measurements of daphnids from culture medium had shown Cd levels to be below MDL. Hence, the initial Cd content of the daphnids could be neglected in the calculations.

Integration of equation (1) between start ([Cd]_daphnia_ =0) and end of the 4-day exposure period (t_exposure_=4d) (measured Cd in A-daphnids, [Cd]_daphnia,A_) gives:

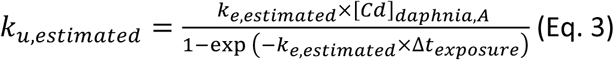

These uptake rates were expressed on a per unit dry body weight basis for data-analyses (Table 1, k_u_*). Raw data of measured Cd in A-daphnids and E-daphnids, body weights, as well as estimated k_u,_ k_u_* and k_e_ for every replicate beaker for every isolate are given in Table 1. Summary data are reported as mean ± standard deviation (n). The data were analyzed using a mixed linear model with ‘Clade’ (‘adapted’ vs non-adapted’ isolates) as a fixed factor, and ‘isolate’ as a random factor nested within ‘Clade’. The dependent variables were uptake rate constant (k_u,estimated_) and elimination rate constant (k_e,estimated_). Statistical analysis was performed in Statistica 7.0 (Statsoft, Tulsa, OK).

## The authors declare no conflicts of interest

## Data Accessibility

The NONA DNA sequencing data is archived and publicly available in the NCBI short read archives under Bioproject ID PRJNA522033. The ADAP genotype DNA sequencing data is archived and publicly available in the NCBI short read archives under Bioproject ID PRJNA1249065.

## Acknowledgements

The authors thank Brandon Mayes and the Cooperative Freshwater Ecology Unit at Laurentian University, who aided in the collection of the Sudbury and Dorset Daphnia isolates used in this study, Jeff Dudycha who generated and supplied the F1 hybrids and parental isolates used in these studies, Haley Waldkoetter, and Elizabeth Turner for assistance on the hybrid vigor and population adaptation studies, and Paul Schumann for valuable help maintaining daphniid cultures and MA lines. We also thank Norman Yan and Bill Keller for helpful discussions on the impact of mining on zooplankton in Sudbury area lakes, and Michael Lynch for discussions on germline mutation, all of which greatly improved this manuscript.

## Author Contributions

N.K. and J.R.S. designed all analyses. N.K., C.J., and J.R.S. analyzed next generation sequencing data. S.G. and K.Y. maintained MA lines. J.R.S. and J.K.C. identified all Daphnia natural populations. K.D. and S.G. designed and implemented biokinetic studies and analyzed the results. All authors contributed edits to the final manuscript.

