## Supplemental File 1 for "Discovery and evaluation of cadmium-adapted *Daphnia pulex* genotypes in a region of historical mining reveals adaptation protects the germline from cadmium-induced mutations"

**Supplemental Material**

**Figure S1**

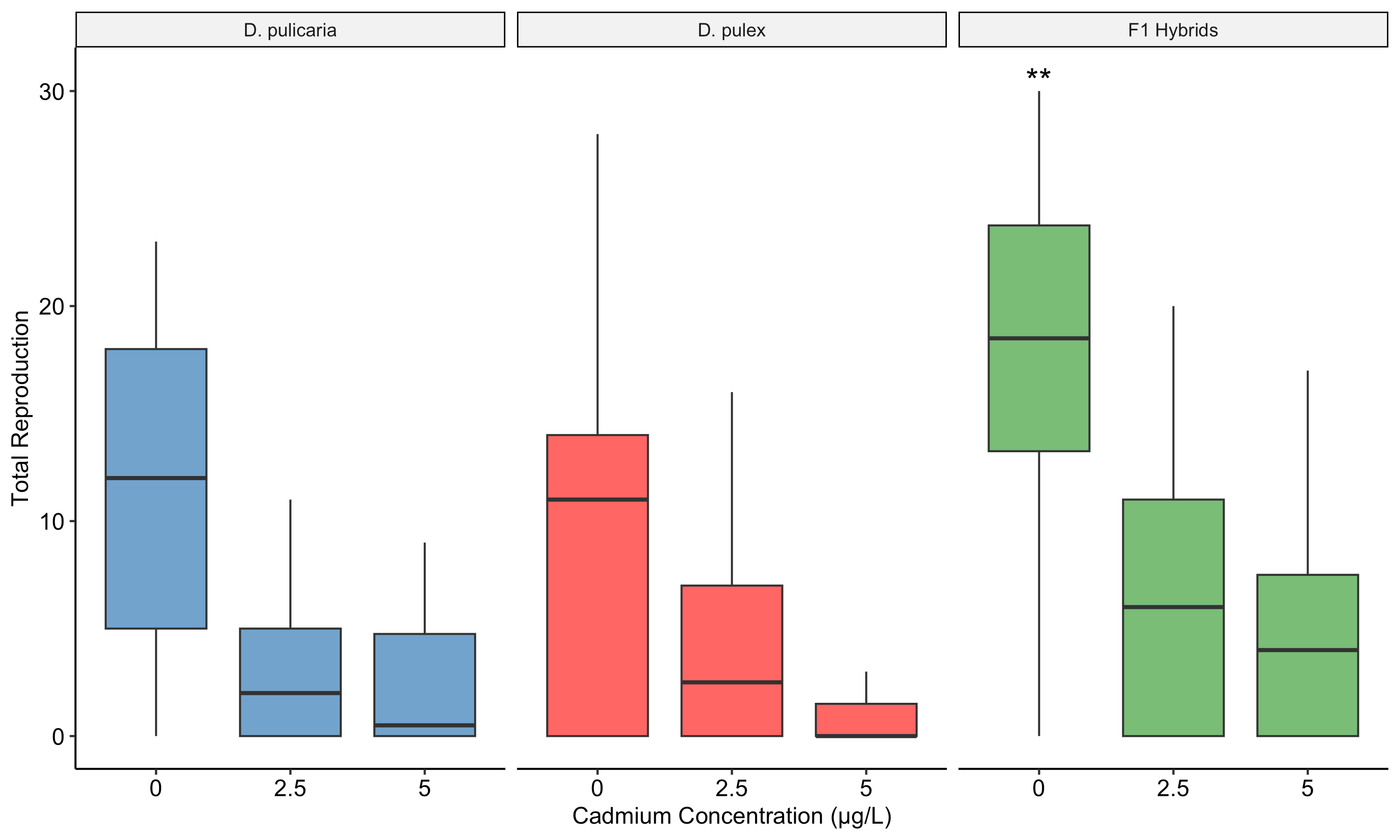

Boxplots show the total reproduction of *D. pulicaria, D. pulex,* and their F_1_ hybrids across exposure to three cadmium concentrations (0, 2.5, and 5 µg/L). Strain was included as a nested random effect in the linear mixed-effect model and each strain consisted of individual biological replicates (n = 5 per strain). Asterisks (**) denote a significant difference within the 0 µg/L conditions between the F_1_ hybrids and the mid-parent mean for the parental strains, indicating hybrid vigor within control conditions (P = 0.006). No significant differences were found between the F_1_ hybrids and either parental strain or mid-parent mean in both cadmium concentrations, indicating that general hybrid vigor in control conditions does not confer a specific, enhanced tolerance to cadmium.

**Figure S2. NONA and ADAP multinucleotide mutations**

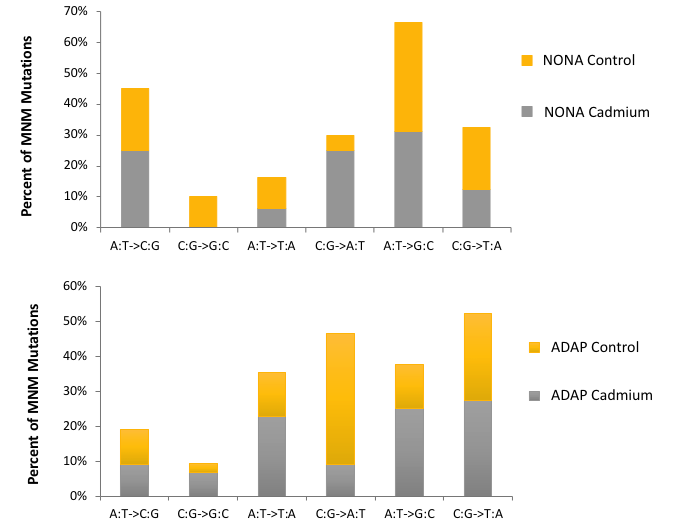

**Figure S3.**

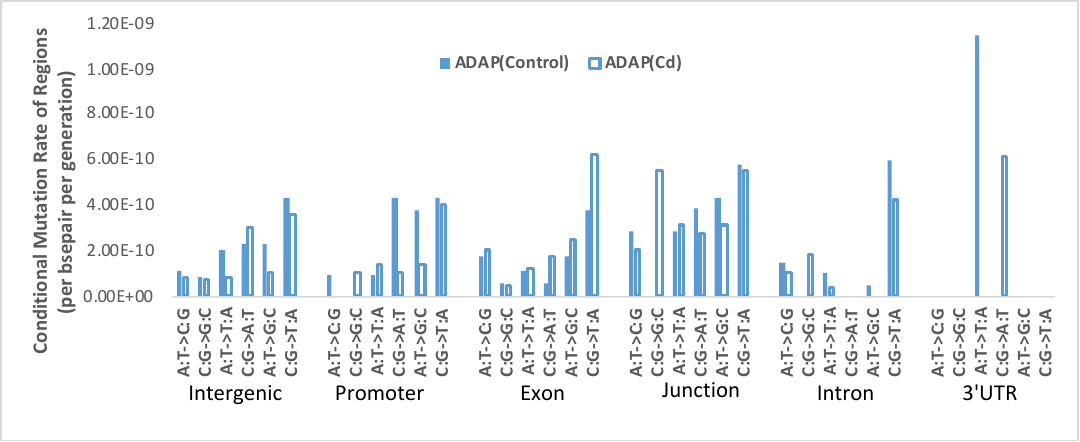

**Figure S4.**

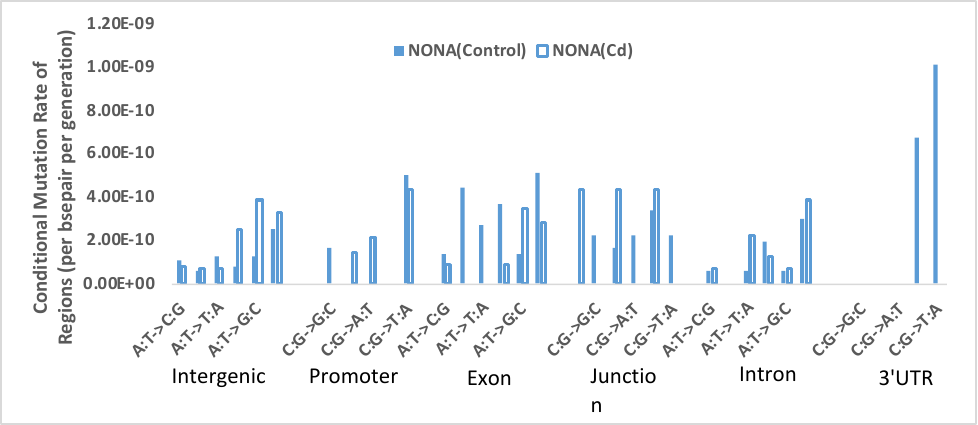

**Figure S5. Total Base Pairs per generation via CNVs (> 3kb)**

Total base pairs per generation

Base Pairs

|  | Total de novo CNV bps per generation |
| --- | --- |
| NONA Control | 818.038 |
| NONA Cadmium | 43.7882 |
| ADAP Control | 8315.79 |
| ADAP Cadmium | 4572.46 |
| LIN | 13111.4 |
| TCO | 3056.86 |

**LIN and TCO (Keith et al. 2016)**

**Figure S6. The number of CNV mutations per generation (CNVs > 3kb)**

de novo CNV mutations per generation

|  | de novo CNVs per generation |
| --- | --- |
| NONA Control | 0.0158228 |
| NONA Cadmium | 0.006 |
| ADAP Control | 0.0944669 |
| ADAP Cadmium | 0.0570048 |
| LIN | 0.144118 |
| TCO | 0.0235294 |

**LIN and TCO (Keith et al. 2016)**

**Supplemental Table Legends**

**Table S1. Lake populations and coordinates sampled from Sudbury and Dorset, Ontario, Canada**

**Table S2. Genome-wide mutation rate data for Non-adapted genotype in control conditions. “**Depth of Coverage” is the genome-wide average depth of sequencing coverage after mapping. Generations is abbreviated as **“**Gens.”. “No. of mutations” is the number of single nucleotide mutations. “Ts/Tv ratio” is the ratio of transitions to transversions. Single nucleotide mutation is abbreviated as “SNM”. “S.E.” is the standard error.

**Table S3. Genome-wide mutation rate data for Adapted genotype in control conditions. “**Depth of Coverage” is the genome-wide average depth of sequencing coverage after mapping. Generations is abbreviated as **“**Gens.”. “No. of mutations” is the number of single nucleotide mutations. “Ts/Tv ratio” is the ratio of transitions to transversions. Single nucleotide mutation is abbreviated as “SNM”. “S.E.” is the standard error.

**Table S4. Conditional mutation rate results for Non-adapted genotype in control conditions**. The overall conditional mutation rate for each mutation class is listed at the bottom of the table, with the S.E. directly below.

**Table S5. Conditional mutation rate results for Adapted genotype in control conditions.**

The overall conditional mutation rate for each mutation class is listed at the bottom of the table, with the S.E. directly below.

**Table S6. Genome-wide mutation rate data for Nonadapted genotype in cadmium exposure. “**Depth of Coverage” is the genome-wide average depth of sequencing coverage after mapping. Generations is abbreviated as **“**Gens.”. “No. of mutations” is the number of single nucleotide mutations. “Ts/Tv ratio” is the ratio of transitions to transversions. Single nucleotide mutation is abbreviated as “SNM”. “S.E.” is the standard error.

**Table S7. Genome-wide mutation rate data for Adapted genotype in cadmium exposure.**

**“**Depth of Coverage” is the genome-wide average depth of sequencing coverage after mapping. Generations is abbreviated as **“**Gens.”. “No. of mutations” is the number of single nucleotide mutations. “Ts/Tv ratio” is the ratio of transitions to transversions. Single nucleotide mutation is abbreviated as “SNM”. “S.E.” is the standard error.

**Table S8. Conditional mutation rate results for Nonadapted genotype in cadmium exposure.** The overall conditional mutation rate for each mutation class is listed at the bottom of the table, with the S.E. directly below.

**Table S9. Conditional mutation rate results for Adapted genotype in cadmium exposure.**

The overall conditional mutation rate for each mutation class is listed at the bottom of the table, with the S.E. directly below.

**Table S10-S13. Context-dependent mutation results for Nonadapted genotype in control conditions.** “Tot. Trips” is the number of observed triplets analyzed in the genome. “Mut Trips” is the number of observed mutations for each type of triplet. Expected is the expectation if mutations were randomly distributed genome-wide. “CpG” refers to all combined contexts where the site of the mutation was originally a C or G, and was flanked on either or both sides by C or G.

**Table S14. Proportion of total mutations in discrete genome regions compared to the random expectation**

**Table S15. Pairwise comparisons of the proportion of total mutations in specific genome regions**

**Table S18. Summary CNV information for NONA and ADAP in both controls and cadmium exposure.**

Summary CNV information for NONA Control, NONA Cadmium, ADAP Control, ADAP Cadmium, and CNV findings from Keith et al. (2016) *D. pulex* experiment are listed. From Keith et al., ASEX is an obligate asexual *D. pulex* genotype, and SEX is a cyclical parthenogen (sexual) *D. pulex* genotype.

**Table S19. Genome Coordinates of CNVs**

Sub-line, Chromosome, Scaffold, First Position, Last Position, and CNV Length are listed. Chromosome coordinates are from the TCO genetic map (Xu et al. 2016). First Position and Last Position are listed in the 5’ to 3’ orientation on the scaffolds where they are observed. CNVs on chromosomes listed as “n/a” are CNVs on scaffolds that were not mapped by Xu et al. (2016) to specific chromosomes.

**Table S16. Summary CNV information for NONA and ADAP in both controls and cadmium exposure**

**Table S17. Genome Coordinates of CNVs**

**Table S1. Lake populations and coordinates sampled from Sudbury and Dorset, Ontario, Canada**

| Lake |  |  | Region |  | Latitude | Longitude |
| --- | --- | --- | --- | --- | --- | --- |
| Basshaunt |  |  | Dorset |  | 45° 7'26.38"N | 78°27'47.19"W |
| Brandy |  |  | Dorset |  | 45° 6'25.74"N | 79°31'35.71"W |
| Buck |  |  | Dorset |  | 45°23'32.55"N | 78°59'29.87"W |
| Crown |  |  | Dorset |  | 45°26'6.54"N | 78°40'7.22"W |
| Frenchman |  |  | Sudbury |  | 46°42'47.78"N | 80°59'6.74"W |
| Glen |  |  | Dorset |  | 45° 7'53.53"N | 78°28'32.14"W |
| Joe |  |  | Sudbury |  | 46°44'5.21"N | 81° 0'42.96"W |
| Kelly |  |  | Sudbury |  | 46°26'48.05"N | 81° 4'0.95"W |
| Leech |  |  | Dorset |  | 45° 3'10.12"N | 79° 5'57.60"W |
| MacFarlane |  |  | Sudbury |  | 46°25'0.79"N | 80°57'43.92"W |
| McCharles |  |  | Sudbury |  | 46°22'55.29"N | 81°14'18.36"W |
| Ramsey |  |  | Sudbury |  | 46°28'34.15"N | 80°58'38.58"W |
| Simon |  |  | Sudbury |  | 46°23'53.95"N | 81°11'18.95"W |

**Table S2. Genome-wide mutation rate data for Nonadapted genotype in control conditions**

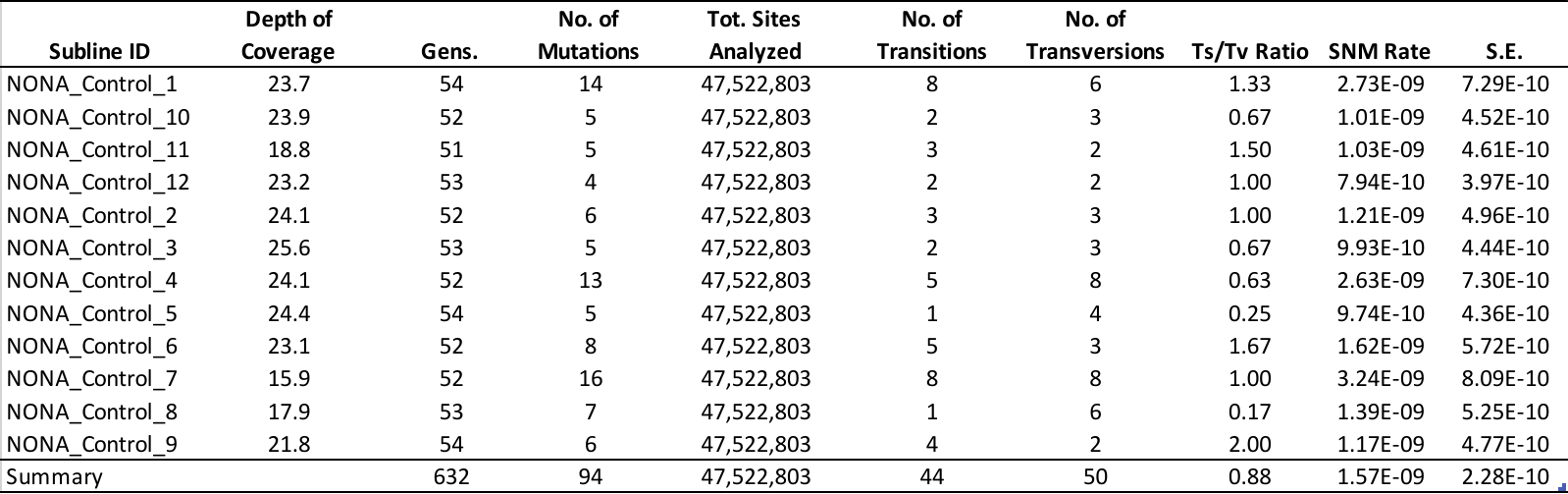

**Table S3. Genome-wide mutation rate data for Adapted genotype in control conditions**

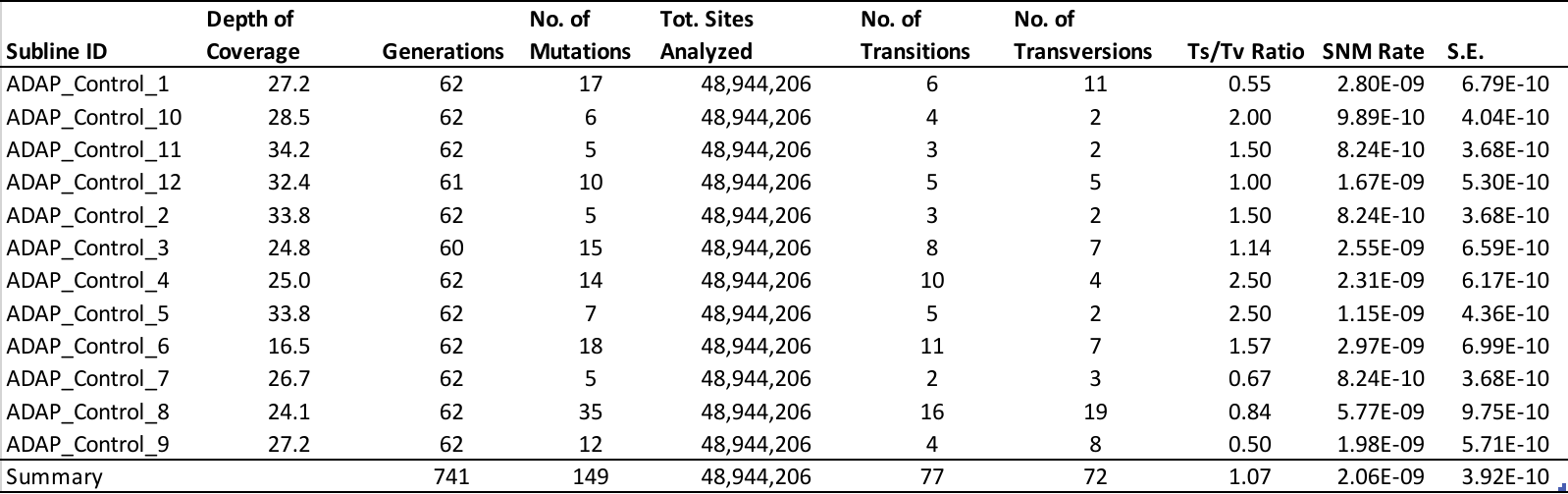

**Table S4. Conditional mutation rate results for Nonadapted genotype in control conditions**

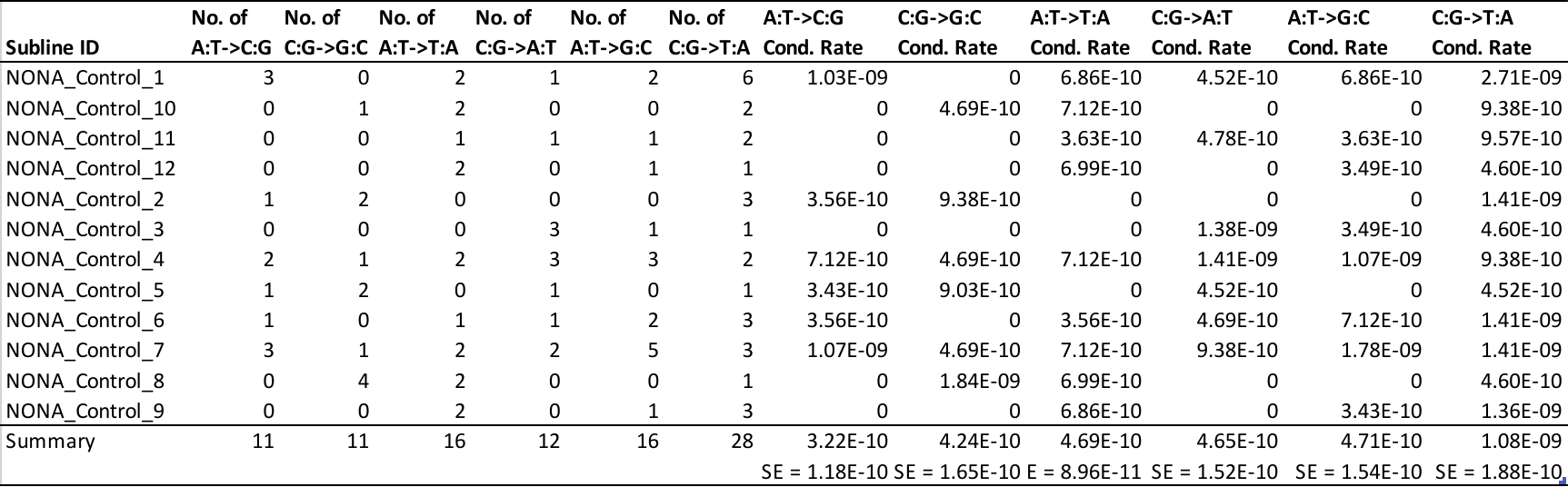

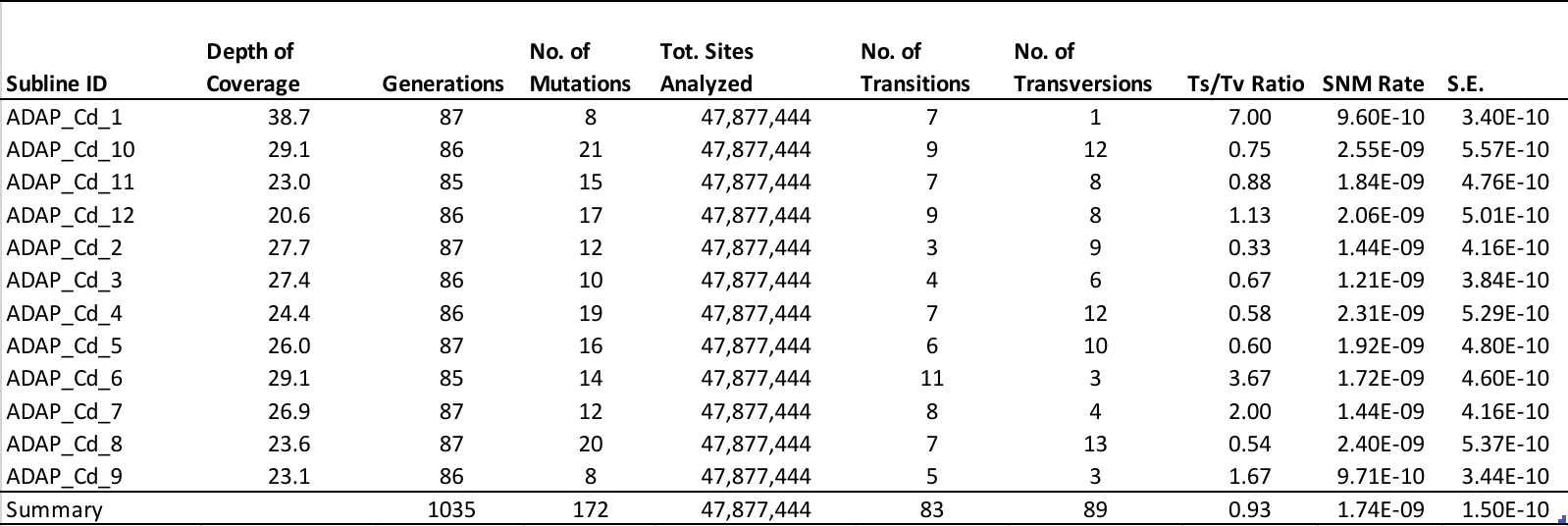

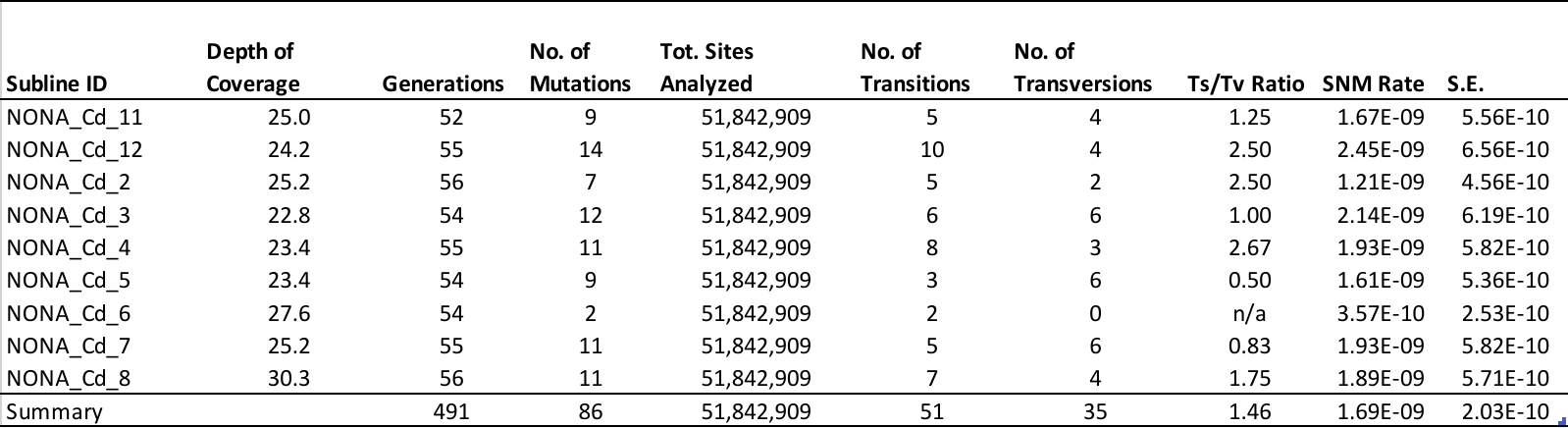

**Table S7. Genome-wide mutation rate data for Adapted genotype in cadmium exposure**

**Table S6. Genome-wide mutation rate data for Nonadapted genotype in cadmium exposure**

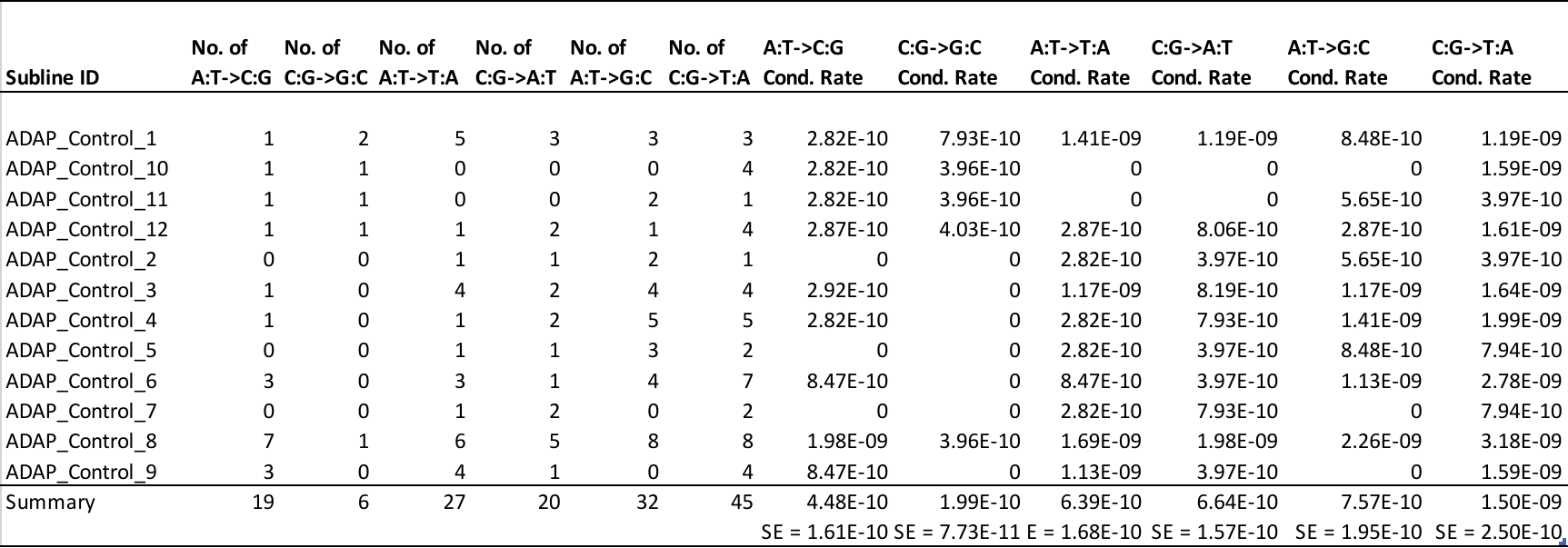

**Table S5. Conditional mutation rate results for Adapted genotype in control conditions**

**Table S9. Conditional mutation rate results for Adapted genotype in cadmium exposure**

**Table S8. Conditional mutation rate results for Nonadapted genotype in cadmium exposure**

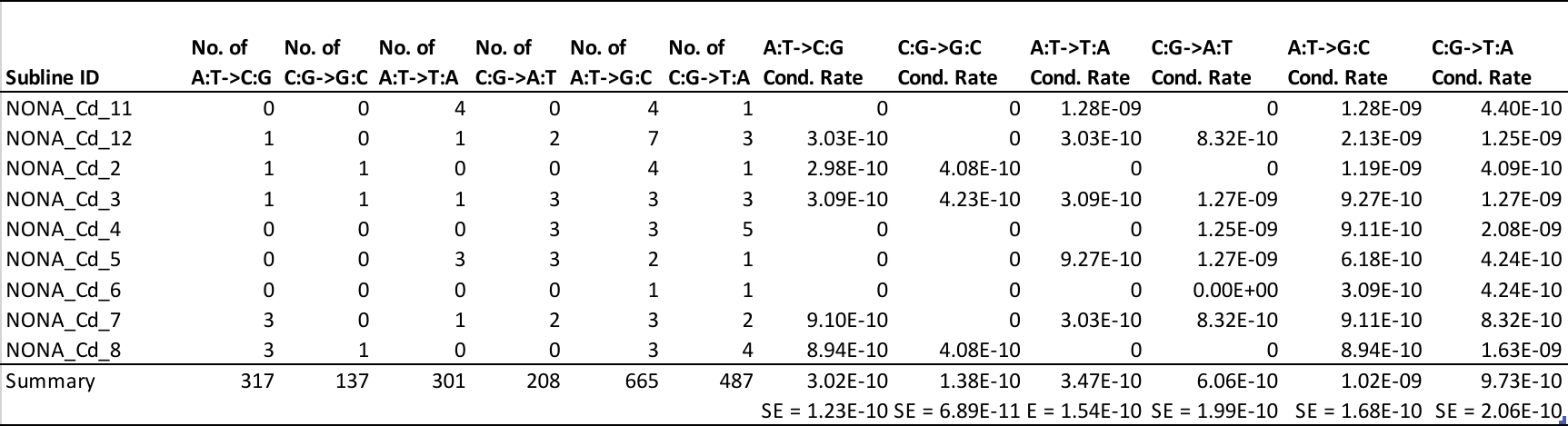

**Table S9. Conditional mutation rate results for Adapted genotype in cadmium exposure**

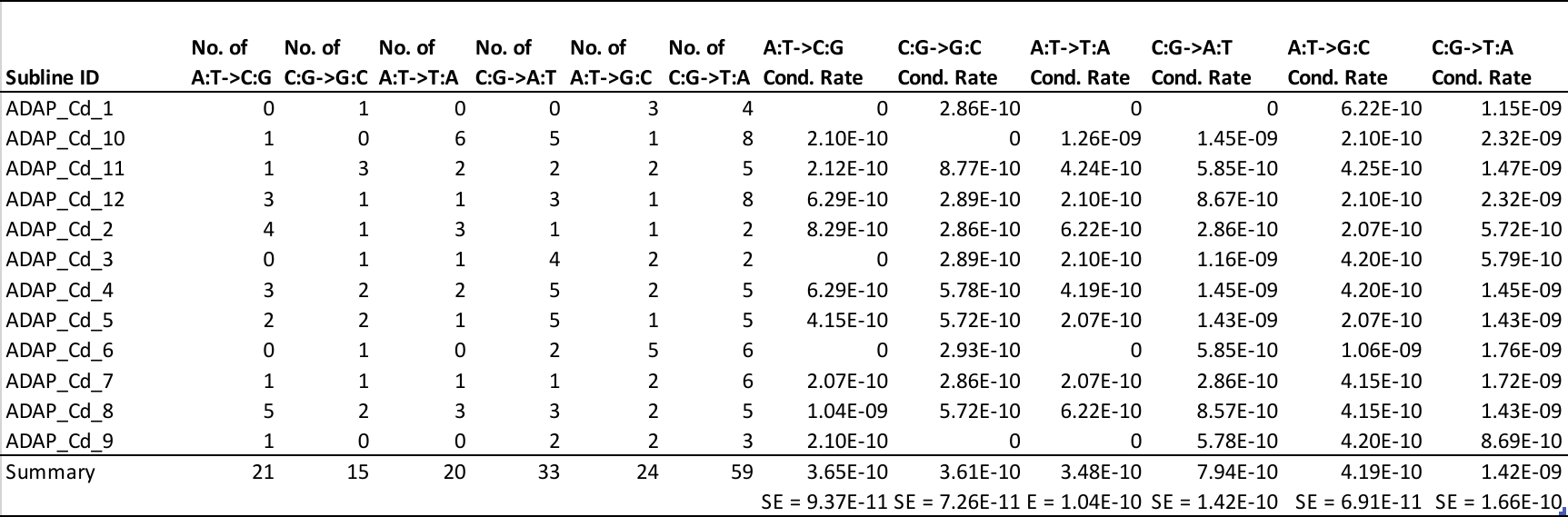

**Table S10. Context-dependent mutation results for Nonadapted genotype in control conditions**

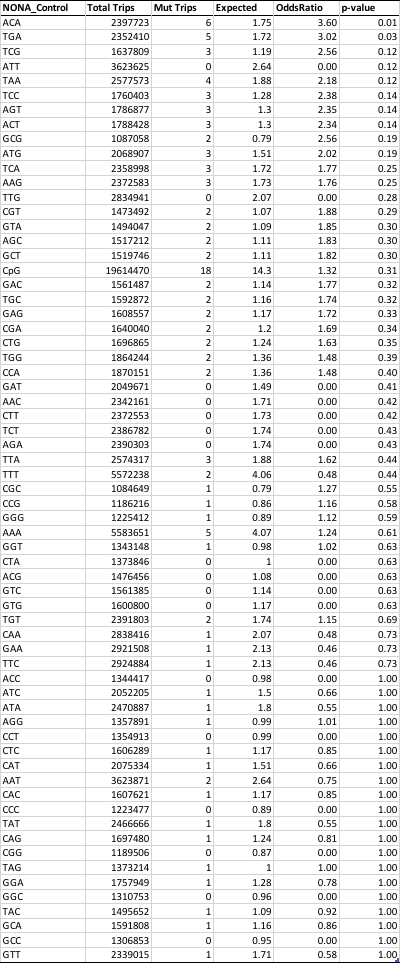

**Table S11. Context-dependent mutation results for Adapted genotype in control conditions**

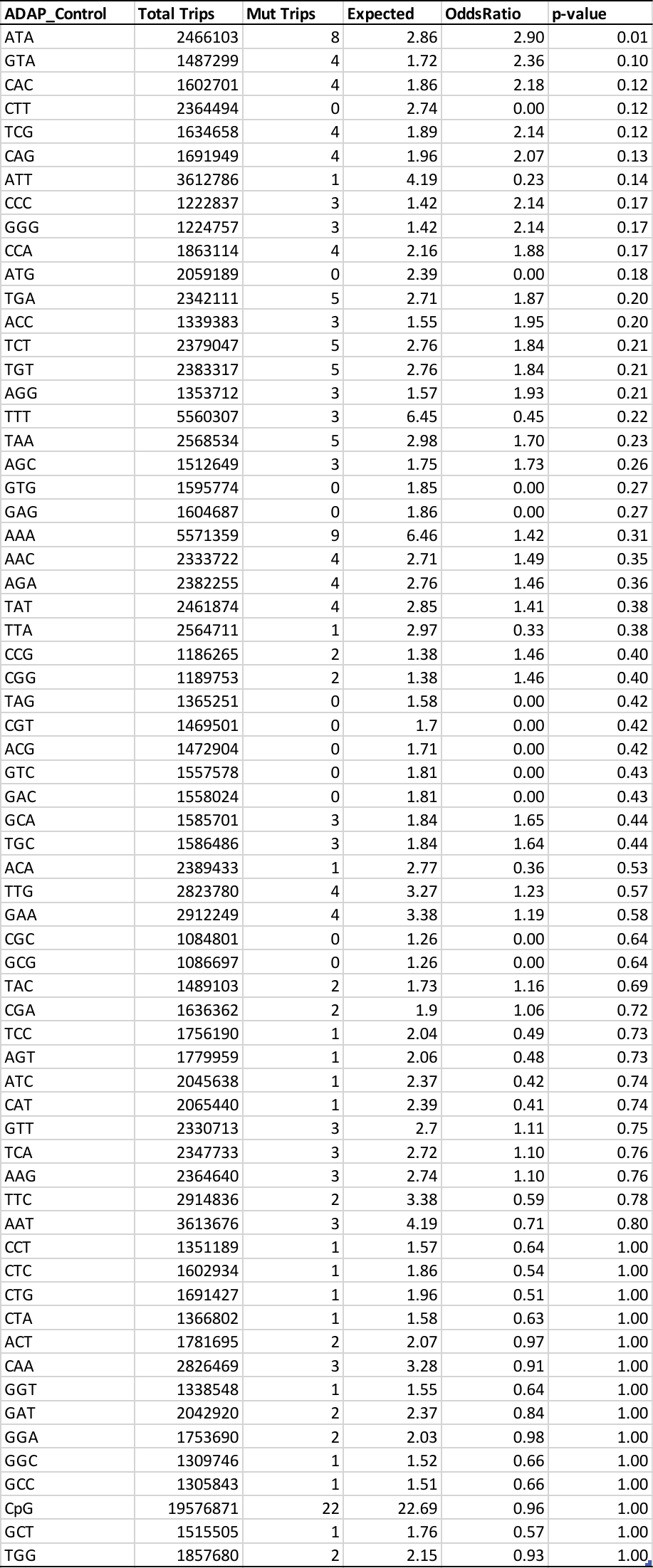

**Table S12. Context-dependent mutation results for Nonadapted genotype in cadmium exposure**

**Table S13. Context-dependent mutation results for Adapted genotype in cadmium exposure**

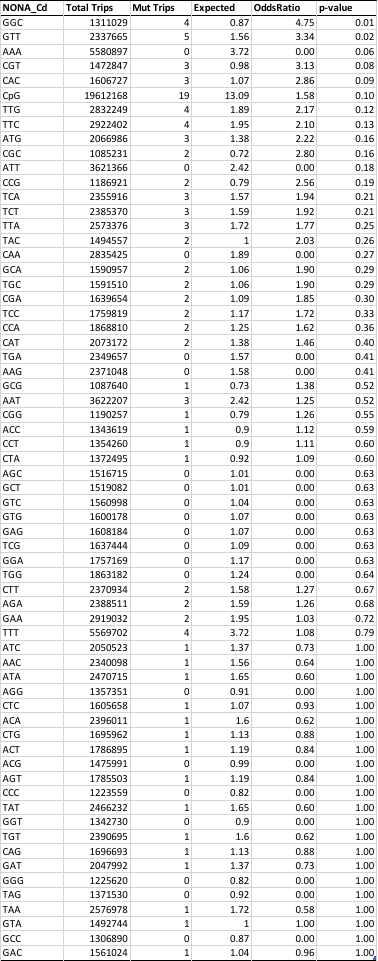

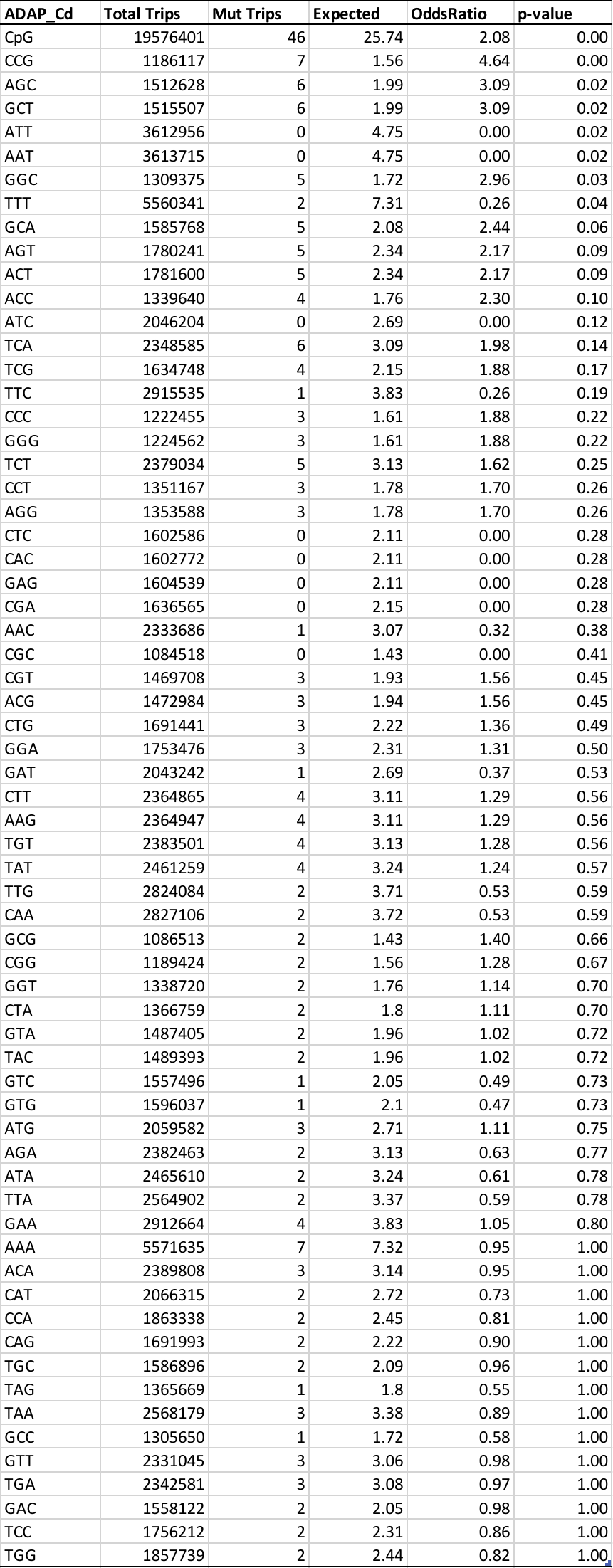

**Table S14. Proportion of total mutations in discrete genome regions compared to the random expectation**

| **Genotype (Condition)** | **Genome Region** | **Proportion of Mutations** | **P-value** |
| --- | --- | --- | --- |
| NONA (Control) | Intergenic | 0.511 | 0.0727 |
| NONA (Control) | Promoter | 0.042 | 0.4179 |
| **NONA (Control)** | **Exon** | **0.277** | **0.0003***** |
| NONA (Control) | Splice Junction | 0.064 | 0.4686 |
| NONA (Control) | Intron | 0.085 | 0.2791 |
| NONA (Control) | 3-UTR | 0.021 | 0.3114 |
| NONA (Cadmium) | Intergenic | 0.674 | 0.1879 |
| NONA (Cadmium) | Promoter | 0.047 | 0.527 |
| NONA (Cadmium) | Exon | 0.105 | 0.5271 |
| NONA (Cadmium) | Splice Junction | 0.07 | 0.318 |
| NONA (Cadmium) | Intron | 0.105 | 0.6291 |
| NONA (Cadmium) | 3-UTR | 0 | 0.6301 |
| ADAP (Control) | Intergenic | 0.631 | 0.5578 |
| ADAP (Control) | Promoters | 0.081 | 0.6307 |
| ADAP (Control) | Exons | 0.107 | 0.4005 |
| ADAP (Control) | Splice Junctions | 0.081 | 0.0845 |
| ADAP (Control) | Introns | 0.087 | 0.1425 |
| ADAP (Control) | 3' UTRs | 0.013 | 0.7005 |
| ADAP (Cadmium) | Intergenic | 0.558 | 0.2421 |
| ADAP (Cadmium) | Promoters | 0.058 | 0.6549 |
| **ADAP (Cadmium)** | **Exons** | **0.192** | **0.0337*** |
| **ADAP (Cadmium)** | **Splice Junctions** | **0.105** | **0.0022**** |
| ADAP (Cadmium) | Introns | 0.081 | 0.0679 |
| ADAP (Cadmium) | 3' UTRs | 0.006 | 0.728 |

**Table S15. Pairwise comparisons of the proportion of total mutations in specific genome regions**

| **Experimental Comparison** | **Genome Region** | **P-value** |
| --- | --- | --- |
| **NONA (Control) vs. NONA (Cd)** | **Intergenic** | **0.0336*** |
|  | Promoters | 1 |
|  | **Exons** | **0.0044**** |
|  | Junctions | 1 |
|  | Introns | 0.7997 |
|  | 3'UTRs | 0.4981 |
| **ADAP (Control) vs. ADAP (Cd)** | Intergenic | 0.2107 |
|  | Promoters | 0.5086 |
|  | **Exons** | **0.0429*** |
|  | Junctions | 0.565 |
|  | Introns | 1 |
|  | 3'UTRs | 0.5987 |
| **NONA (Control) vs. ADAP (Control)** | Intergenic | 0.0821 |
|  | Promoters | 0.2970 |
|  | **Exons** | **0.0009***** |
|  | Junctions | 0.8025 |
|  | Introns | 1 |
|  | 3'UTRs | 0.6416 |
| **NONA (Cd) vs. ADAP (Cd)** | Intergenic | 0.0809 |
|  | Promoters | 0.7796 |
|  | Exons | 0.0769 |
|  | Junctions | 0.4960 |
|  | Introns | 0.6435 |
|  | 3'UTRs | 1 |
| **NONA (Control) vs. ADAP (Cd)** | Intergenic | 0.5201 |
|  | Promoters | 0.7759 |
|  | Exons | 0.124 |
|  | Junctions | 0.371 |
|  | Introns | 1 |
|  | 3'UTRs | 0.2856 |
| **NONA (Cd) vs. ADAP (Control)** | Intergenic | 0.5714 |
|  | Promoters | 0.4241 |
|  | Exons | 1 |
|  | Junctions | 1 |
|  | Introns | 0.6496 |
|  | 3'UTRs | 0.5339 |

Table 16. 5-hmC readings for ADAP and NONA at concentrations 0, 0.25. and 20 μg Cd /L

|  | **ADAP 0** | **ADAP 0.25** | **ADAP 20** | **NONA 0** | **NONA 0.25** | **NONA 20** |
| --- | --- | --- | --- | --- | --- | --- |
| **Rep 1** | 0.259 | 0.249 | 0.260 | debris in well | 0.414 | 0.296 |
| **Rep 2** | 0.258 | 0.165 | 0.151 | 0.679 | 0.140 | 0.424 |
| **Rep 3** | 0.178 | 0.317 | 0.381 | 0.434 | 0.167 | 0.248 |
| **Rep 4** | 0.258 | 0.315 | 0.414 | 0.477 | 0.115 | 0.270 |
| **Avg** | 0.238 | 0.244 | 0.301 | 0.530 | 0.209 | 0.310 |
| **Std Dev** | 0.040 | 0.076 | 0.120 | 0.131 | 0.138 | 0.079 |

**Table S17. Summary CNV information for NONA and ADAP in both controls and cadmium exposure**

| **Genotype/Condition** | **No. of CNVs** | **Med. Length** | **Avg. Length** | **Total CNV BPs** | **CNV BPs / gen.** | **CNV events / gen.** |
| --- | --- | --- | --- | --- | --- | --- |
| NONA Control | 10 | 29,750 | 51,700 | 517,000 | 818 | 0.016 |
| NONA Cadmium | 3 | 4,000 | 7,167 | 21,500 | 44 | 0.006 |
| ADAP Control | 70 | 11,500 | 88,029 | 6,161,999 | 8,316 | 0.095 |
| ADAP Cadmium | 59 | 8,500 | 80,212 | 4,732,500 | 4,572 | 0.057 |
| ASEX - Keith et al. (2016) | 98 | 17,000 | 90,977 | 8,915,750 | 13,111 | 0.144 |
| SEX - Keith et al. (2016) | 6 | 30,250 | 129,917 | 779,500 | 3,056 | 0.024 |

**Table S18. Genome Coordinates of CNVs**

| **Subline** | **Chromosome** | **Scaffold** | **First Position** | **Last Position** | **CNV length** |
| --- | --- | --- | --- | --- | --- |
| NONA Control 2 | 4 | 43 | 803500 | 842500 | 39000 |
| NONA Control 12 | 5 | 131 | 0 | 212500 | 212500 |
| NONA Control 12 | 5 | 131 | 235500 | 274500 | 39000 |
| NONA Control 12 | 5 | 39 | 272000 | 309500 | 37500 |
| NONA Control 12 | 5 | 89 | 5500 | 20000 | 14500 |
| NONA Control 12 | 6 | 47 | 911500 | 916500 | 5000 |
| NONA Control 1 | 8 | 22 | 1177500 | 1181000 | 3500 |
| NONA Control 12 | 8 | 20 | 351000 | 489500 | 138500 |
| NONA Control 3 | 9 | 9 | 620500 | 642500 | 22000 |
| NONA Cd 1 | n/a | 69 | 572000 | 577500 | 5500 |
| NONA Cd 3 | 2 | 1 | 3116500 | 3130000 | 13500 |
| NONA Cd 5 | 11 | 67 | 165500 | 169500 | 4000 |
| NONA Cd 5 | 11 | 67 | 261500 | 265500 | 4000 |
| ADAP Control 9 | 2 | 1 | 32000 | 128000 | 96000 |
| ADAP Control 9 | 2 | 1 | 3243000 | 3751500 | 508500 |
| ADAP Control 9 | 2 | 19 | 86000 | 247500 | 161500 |
| ADAP Control 11 | 2 | 19 | 704500 | 1304000 | 599500 |
| ADAP Control 2 | 3 | 2 | 3093000 | 3096500 | 3500 |
| ADAP Control 10 | 3 | 16 | 518000 | 525500 | 7500 |
| ADAP Control 10 | 3 | 21 | 529500 | 590500 | 61000 |
| ADAP Control 10 | 3 | 21 | 681500 | 688500 | 7000 |
| ADAP Control 10 | 3 | 44 | 6500 | 979500 | 973000 |
| ADAP Control 10 | 3 | 114 | 9000 | 114500 | 105500 |
| ADAP Control 10 | 3 | 114 | 147500 | 347000 | 199500 |
| ADAP Control 10 | 3 | 155 | 54500 | 125000 | 70500 |
| ADAP Control 10 | 3 | 164 | 23000 | 64500 | 41500 |
| ADAP Control 10 | 3 | 164 | 122000 | 127000 | 5000 |
| ADAP Control 10 | 3 | 164 | 147000 | 153000 | 6000 |
| ADAP Control 10 | 3 | 164 | 239500 | 258000 | 18500 |
| ADAP Control 10 | 3 | 174 | 141000 | 145500 | 4500 |
| ADAP Control 10 | 3 | 174 | 166500 | 201500 | 35000 |
| ADAP Control 10 | 3 | 174 | 222000 | 235000 | 13000 |
| ADAP Control 10 | 3 | 178 | 43500 | 58000 | 14500 |
| ADAP Control 10 | 3 | 178 | 85500 | 89500 | 4000 |
| ADAP Control 10 | 3 | 178 | 141500 | 188500 | 47000 |
| ADAP Control 10 | 5 | 15 | 1364000 | 1380500 | 16500 |
| ADAP Control 7 | 5 | 60 | 1 | 617500 | 617499 |
| ADAP Control 1 | 6 | 32 | 82500 | 381500 | 299000 |
| ADAP Control 11 | 6 | 32 | 404000 | 423500 | 19500 |
| ADAP Control 7 | 6 | 32 | 424000 | 428000 | 4000 |
| ADAP Control 10 | 7 | 40 | 42000 | 45000 | 3000 |
| ADAP Control 3 | 7 | 48 | 454500 | 457500 | 3000 |
| ADAP Control 10 | 7 | 82 | 62500 | 69000 | 6500 |
| ADAP Control 10 | 7 | 87 | 386500 | 389500 | 3000 |
| ADAP Control 5 | 8 | 34 | 528500 | 531500 | 3000 |
| ADAP Control 10 | 8 | 57 | 291500 | 296000 | 4500 |
| ADAP Control 6 | 9 | 9 | 1396000 | 1410500 | 14500 |
| ADAP Control 11 | 9 | 41 | 465000 | 470500 | 5500 |
| ADAP Control 10 | 9 | 107 | 4500 | 8000 | 3500 |
| ADAP Control 10 | 10 | 6 | 1873500 | 1888500 | 15000 |
| ADAP Control 10 | 10 | 17 | 0 | 454500 | 454500 |
| ADAP Control 12 | 10 | 26 | 740500 | 853000 | 112500 |
| ADAP Control 12 | 10 | 26 | 1002000 | 1013000 | 11000 |
| ADAP Control 10 | 10 | 72 | 19000 | 384500 | 365500 |
| ADAP Control 5 | 10 | 72 | 396000 | 409000 | 13000 |
| ADAP Control 5 | 10 | 72 | 449500 | 459500 | 10000 |
| ADAP Control 10 | 10 | 72 | 573000 | 642500 | 69500 |
| ADAP Control 10 | 10 | 173 | 43500 | 237500 | 194000 |
| ADAP Control 6 | 11 | 8 | 413000 | 497500 | 84500 |
| ADAP Control 4 | 11 | 10 | 2061000 | 2153500 | 92500 |
| ADAP Control 4 | 11 | 67 | 12500 | 15500 | 3000 |
| ADAP Control 4 | 11 | 67 | 53500 | 57000 | 3500 |
| ADAP Control 4 | 11 | 67 | 77500 | 113000 | 35500 |
| ADAP Control 4 | 11 | 67 | 139000 | 158500 | 19500 |
| ADAP Control 4 | 11 | 67 | 367500 | 713000 | 345500 |
| ADAP Control 9 | 11 | 67 | 367500 | 370500 | 3000 |
| ADAP Control 9 | 11 | 67 | 480500 | 690000 | 209500 |
| ADAP Control 10 | n/a | 50 | 118500 | 122000 | 3500 |
| ADAP Control 10 | n/a | 122 | 255500 | 265000 | 9500 |
| ADAP Control 10 | n/a | 122 | 288500 | 292000 | 3500 |
| ADAP Control 10 | n/a | 122 | 312500 | 323500 | 11000 |
| ADAP Control 10 | n/a | 122 | 371500 | 379000 | 7500 |
| ADAP Control 10 | n/a | 132 | 140000 | 144500 | 4500 |
| ADAP Control 10 | n/a | 132 | 175000 | 183500 | 8500 |
| ADAP Control 10 | n/a | 132 | 211500 | 215000 | 3500 |
| ADAP Control 10 | n/a | 132 | 279000 | 282000 | 3000 |
| ADAP Control 10 | n/a | 147 | 275500 | 287500 | 12000 |
| ADAP Control 10 | n/a | 151 | 125500 | 130000 | 4500 |
| ADAP Control 10 | n/a | 168 | 93000 | 145000 | 52000 |
| ADAP Control 10 | n/a | 168 | 256000 | 261000 | 5000 |
| ADAP Control 10 | n/a | 196 | 32000 | 36000 | 4000 |
| ADAP Cd 6 | 1 | 3 | 3679500 | 3777500 | 98000 |
| ADAP Cd 6 | 1 | 53 | 0 | 504000 | 504000 |
| ADAP Cd 2 | 1 | 66 | 189000 | 221500 | 32500 |
| ADAP Cd 6 | 1 | 66 | 590500 | 807500 | 217000 |
| ADAP Cd 6 | 1 | 106 | 0 | 493500 | 493500 |
| ADAP Cd 2 | 1 | 130 | 116500 | 125000 | 8500 |
| ADAP Cd 6 | 1 | 130 | 139500 | 326000 | 186500 |
| ADAP Cd 4 | 2 | 78 | 299000 | 520500 | 221500 |
| ADAP Cd 4 | 3 | 2 | 966500 | 971000 | 4500 |
| ADAP Cd 2 | 3 | 189 | 112000 | 116000 | 4000 |
| ADAP Cd 11 | 4 | 31 | 497000 | 504000 | 7000 |
| ADAP Cd 11 | 4 | 31 | 543000 | 550500 | 7500 |
| ADAP Cd 11 | 4 | 31 | 577500 | 581000 | 3500 |
| ADAP Cd 11 | 4 | 31 | 594500 | 598500 | 4000 |
| ADAP Cd 11 | 4 | 31 | 1016500 | 1025000 | 8500 |
| ADAP Cd 3 | 5 | 15 | 1378000 | 1382500 | 4500 |
| ADAP Cd 2 | 5 | 37 | 90000 | 93000 | 3000 |
| ADAP Cd 12 | 6 | 32 | 252000 | 446500 | 194500 |
| ADAP Cd 7 | 6 | 32 | 409000 | 412500 | 3500 |
| ADAP Cd 10 | 6 | 92 | 203500 | 206500 | 3000 |
| ADAP Cd 10 | 6 | 92 | 476500 | 501000 | 24500 |
| ADAP Cd 12 | 7 | 4 | 963500 | 1019000 | 55500 |
| ADAP Cd 12 | 7 | 4 | 1140000 | 1162000 | 22000 |
| ADAP Cd 4 | 7 | 18 | 1387500 | 1391000 | 3500 |
| ADAP Cd 8 | 7 | 54 | 626500 | 629500 | 3000 |
| ADAP Cd 7 | 7 | 91 | 110000 | 115500 | 5500 |
| ADAP Cd 10 | 8 | 20 | 215500 | 466000 | 250500 |
| ADAP Cd 4 | 8 | 169 | 109500 | 112500 | 3000 |
| ADAP Cd 4 | 8 | 183 | 41500 | 47500 | 6000 |
| ADAP Cd 1 | 8 | 183 | 58000 | 124000 | 66000 |
| ADAP Cd 1 | 8 | 183 | 143000 | 149500 | 6500 |
| ADAP Cd 11 | 9 | 9 | 264000 | 289000 | 25000 |
| ADAP Cd 7 | 9 | 9 | 477000 | 694500 | 217500 |
| ADAP Cd 6 | 9 | 9 | 1104000 | 1108500 | 4500 |
| ADAP Cd 11 | 9 | 9 | 1140500 | 1269000 | 128500 |
| ADAP Cd 4 | 9 | 9 | 1278500 | 1282000 | 3500 |
| ADAP Cd 11 | 9 | 9 | 1281500 | 1312000 | 30500 |
| ADAP Cd 11 | 9 | 9 | 1329000 | 1374500 | 45500 |
| ADAP Cd 11 | 9 | 14 | 1439500 | 1455500 | 16000 |
| ADAP Cd 7 | 10 | 6 | 2290500 | 2297000 | 6500 |
| ADAP Cd 4 | 10 | 17 | 1518000 | 1522500 | 4500 |
| ADAP Cd 10 | 10 | 36 | 931000 | 934000 | 3000 |
| ADAP Cd 10 | 10 | 49 | 10500 | 16500 | 6000 |
| ADAP Cd 6 | 10 | 56 | 114000 | 496500 | 382500 |
| ADAP Cd 2 | 10 | 72 | 390000 | 573000 | 183000 |
| ADAP Cd 4 | 10 | 72 | 452500 | 455500 | 3000 |
| ADAP Cd 4 | 10 | 72 | 573000 | 590500 | 17500 |
| ADAP Cd 2 | 10 | 72 | 625500 | 628500 | 3000 |
| ADAP Cd 6 | 10 | 103 | 57500 | 60500 | 3000 |
| ADAP Cd 6 | 10 | 103 | 271500 | 275000 | 3500 |
| ADAP Cd 1 | 11 | 10 | 1345000 | 2146000 | 801000 |
| ADAP Cd 12 | 11 | 67 | 367000 | 370000 | 3000 |
| ADAP Cd 5 | 11 | 67 | 436000 | 470500 | 34500 |
| ADAP Cd 12 | 11 | 67 | 489500 | 713000 | 223500 |
| ADAP Cd 3 | 12 | 58 | 710000 | 724500 | 14500 |
| ADAP Cd 10 | 12 | 70 | 109500 | 194000 | 84500 |
| ADAP Cd 10 | 12 | 96 | 335500 | 351500 | 16000 |
| ADAP Cd 1 | n/a | 85 | 295000 | 298000 | 3000 |
| ADAP Cd 10 | n/a | 167 | 47500 | 58500 | 11000 |

Table S19. qPCR Results for genomic DNA (gDNA Gene Amplicon Copies) and RNA (RNA Expression qPCR)

| **Group** | **Genotype ID** | **Ct (dRN)** | **Copies** |
| --- | --- | --- | --- |
| Adapted | K10 | 20.62 | 18,070.00 |
| Adapted | K10 | 20.88 | 15,070.00 |
| Adapted | K10 | 20.94 | 14,450.00 |
| Adapted | K10 | 20.95 | 14,370.00 |
| Adapted | K10 | 21.11 | 12,860.00 |
| Adapted | K10 | 21.12 | 12,750.00 |
| Adapted | K2 | 18.51 | 76,930.00 |
| Adapted | K2 | 18.51 | 76,690.00 |
| Adapted | K2 | 18.57 | 73,640.00 |
| Adapted | K2 | 18.59 | 72,700.00 |
| Adapted | K2 | 18.65 | 69,640.00 |
| Adapted | K2 | 18.66 | 69,230.00 |
| Adapted | K3 | 21.03 | 13,610.00 |
| Adapted | K3 | 21.10 | 12,930.00 |
| Adapted | K3 | 21.16 | 12,420.00 |
| Adapted | K3 | 21.33 | 11,050.00 |
| Adapted | K3 | 21.35 | 10,890.00 |
| Adapted | K3 | 21.44 | 10,300.00 |
| Adapted | K9 | 18.60 | 72,410.00 |
| Adapted | K9 | 18.62 | 71,360.00 |
| Adapted | K9 | 18.68 | 68,500.00 |
| Adapted | K9 | 18.85 | 60,800.00 |
| Adapted | K9 | 18.86 | 60,300.00 |
| Adapted | K9 | 18.97 | 56,070.00 |
| Adapted | MC13 | 18.60 | 72,290.00 |
| Adapted | MC13 | 18.62 | 71,360.00 |
| Adapted | MC13 | 18.63 | 70,810.00 |
| Adapted | MC13 | 18.63 | 70,840.00 |
| Adapted | MC14 | 18.15 | 85,300.00 |
| Adapted | MC14 | 18.21 | 81,460.00 |
| Adapted | MC14 | 18.22 | 80,950.00 |
| Adapted | MC14 | 18.26 | 78,680.00 |
| Adapted | MC14 | 18.33 | 75,400.00 |
| Adapted | MC14 | 18.47 | 68,080.00 |
| Adapted | MC8 | 18.57 | 73,750.00 |
| Adapted | MC8 | 18.58 | 72,980.00 |
| Adapted | MC8 | 18.63 | 70,580.00 |
| Adapted | MC8 | 18.63 | 70,840.00 |
| Adapted | MC8 | 18.66 | 69,280.00 |
| Adapted | MF4 | 18.13 | 86,450.00 |
| Adapted | MF4 | 18.17 | 83,800.00 |
| Adapted | MF4 | 18.30 | 76,490.00 |
| Adapted | MF4 | 18.31 | 75,990.00 |
| Adapted | MF4 | 18.41 | 71,220.00 |
| Adapted | MF4 | 18.55 | 64,380.00 |
| Adapted | MF6 | 18.48 | 67,860.00 |
| Adapted | MF6 | 18.49 | 67,270.00 |
| Adapted | MF6 | 18.50 | 66,690.00 |
| Adapted | MF6 | 18.51 | 66,370.00 |
| Adapted | MF6 | 18.52 | 66,110.00 |
| Adapted | S1 | 18.06 | 90,260.00 |
| Adapted | S1 | 18.09 | 88,750.00 |
| Adapted | S1 | 18.14 | 85,710.00 |
| Adapted | S1 | 18.15 | 85,350.00 |
| Adapted | S1 | 18.17 | 83,770.00 |
| Adapted | S14 | 17.94 | 98,330.00 |
| Adapted | S14 | 17.96 | 97,290.00 |
| Adapted | S14 | 18.11 | 87,210.00 |
| Adapted | S9 | 19.49 | 33,800.00 |
| Adapted | S9 | 19.51 | 33,370.00 |
| Adapted | S9 | 19.59 | 31,500.00 |
| Adapted | S9 | 19.61 | 31,070.00 |
| Adapted | S9 | 19.66 | 30,020.00 |
| Adapted | S9 | 19.66 | 29,970.00 |
| Non-Adapted | BH14 | 21.97 | 7,120.00 |
| Non-Adapted | BH14 | 22.03 | 6,841.00 |
| Non-Adapted | BH14 | 22.04 | 6,814.00 |
| Non-Adapted | BH14 | 22.06 | 6,695.00 |
| Non-Adapted | BH14 | 22.25 | 5,886.00 |
| Non-Adapted | BH14 | 22.52 | 4,889.00 |
| Non-Adapted | BH15 | 18.81 | 62,480.00 |
| Non-Adapted | BH15 | 18.83 | 61,520.00 |
| Non-Adapted | BH15 | 18.92 | 58,050.00 |
| Non-Adapted | BH15 | 19.01 | 54,590.00 |
| Non-Adapted | BH15 | 19.06 | 52,460.00 |
| Non-Adapted | BH3 | 18.83 | 61,780.00 |
| Non-Adapted | BH3 | 18.89 | 59,260.00 |
| Non-Adapted | BH3 | 18.91 | 58,540.00 |
| Non-Adapted | BH3 | 18.97 | 56,140.00 |
| Non-Adapted | BH3 | 18.98 | 55,770.00 |
| Non-Adapted | BH3 | 19.00 | 55,030.00 |
| Non-Adapted | BR1 | 19.60 | 36,280.00 |
| Non-Adapted | BR1 | 19.70 | 33,930.00 |
| Non-Adapted | BR1 | 19.70 | 33,950.00 |
| Non-Adapted | BR1 | 19.82 | 31,310.00 |
| Non-Adapted | BR1 | 19.85 | 30,690.00 |
| Non-Adapted | BR16 | 19.59 | 36,500.00 |
| Non-Adapted | BR16 | 19.60 | 36,280.00 |
| Non-Adapted | BR16 | 19.65 | 35,200.00 |
| Non-Adapted | BR16 | 19.66 | 34,760.00 |
| Non-Adapted | BR16 | 19.68 | 34,440.00 |
| Non-Adapted | BR16 | 19.80 | 31,770.00 |
| Non-Adapted | BU12 | 19.16 | 49,300.00 |
| Non-Adapted | BU12 | 19.22 | 47,180.00 |
| Non-Adapted | BU12 | 19.22 | 47,200.00 |
| Non-Adapted | BU12 | 19.25 | 46,250.00 |
| Non-Adapted | BU12 | 19.32 | 44,090.00 |
| Non-Adapted | BU12 | 19.45 | 40,290.00 |
| Non-Adapted | F10 | 18.16 | 84,290.00 |
| Non-Adapted | F10 | 18.22 | 81,140.00 |
| Non-Adapted | F10 | 18.25 | 79,290.00 |
| Non-Adapted | F10 | 18.26 | 78,880.00 |
| Non-Adapted | F10 | 18.29 | 77,420.00 |
| Non-Adapted | F10 | 18.29 | 77,160.00 |
| Non-Adapted | J3 | 18.11 | 87,720.00 |
| Non-Adapted | J3 | 18.11 | 87,540.00 |
| Non-Adapted | J3 | 18.13 | 86,140.00 |
| Non-Adapted | J3 | 18.36 | 73,460.00 |
| Non-Adapted | J3 | 18.41 | 71,090.00 |
| Non-Adapted | J4 | 18.45 | 68,970.00 |
| Non-Adapted | J4 | 18.51 | 66,310.00 |
| Non-Adapted | J4 | 18.65 | 60,290.00 |
| Non-Adapted | J4 | 18.82 | 53,520.00 |
| Non-Adapted | J4 | 18.85 | 52,680.00 |
| Non-Adapted | G11 | 22.10 | 5,576.00 |
| Non-Adapted | G11 | 22.22 | 5,132.00 |
| Non-Adapted | G11 | 22.34 | 4,737.00 |
| Non-Adapted | G11 | 22.54 | 4,105.00 |
| Non-Adapted | G11 | 23.00 | 3,006.00 |
| Non-Adapted | G11 | 23.39 | 2,297.00 |
| Non-Adapted | L7 | 18.29 | 77,470.00 |
| Non-Adapted | L7 | 18.34 | 74,910.00 |
| Non-Adapted | L7 | 18.36 | 73,430.00 |
| Non-Adapted | L7 | 18.45 | 69,010.00 |
| Non-Adapted | L7 | 18.45 | 69,080.00 |
| Non-Adapted | L7 | 18.48 | 67,680.00 |
| Non-Adapted | R15 | 22.87 | 3,288.00 |
| Non-Adapted | R15 | 22.91 | 3,183.00 |
| Non-Adapted | R15 | 22.93 | 3,144.00 |
| Non-Adapted | R15 | 23.01 | 2,987.00 |
| Non-Adapted | R15 | 23.09 | 2,810.00 |
| Non-Adapted | TCO | 18.37 | 73,240.00 |
| Non-Adapted | TCO | 18.42 | 70,830.00 |
| Non-Adapted | TCO | 18.46 | 68,830.00 |

Table S20. RT-qPCR Results for RNA expression – Adapted vs. Nonadapted

| **Genotype** | Class | **0 Cd** | **20 Cd** |
| --- | --- | --- | --- |
| K3 | Adapted | 2.9 | 20.67 |
| K3 | Adapted | 2.87 | 22.66 |
| K3 | Adapted | 3.6 | 22.8 |
| K7 | Adapted | 7.03 | 8.35 |
| K7 | Adapted | 6.97 | 7.98 |
| K7 | Adapted | 7.54 | 8.09 |
| K13 | Adapted | 7.41 | 7 |
| K13 | Adapted | 7.61 | 6.99 |
| K13 | Adapted | 6.15 | 7.87 |
| MC8 | Adapted | 8 | 8.93 |
| MC8 | Adapted | 8.35 | 10.08 |
| MC8 | Adapted | 8.62 | 8.63 |
| MF6 | Adapted | 7.58 | 16.44 |
| MF6 | Adapted | 8.41 | 15.79 |
| MF6 | Adapted | 9.19 | 15.59 |
| S9 | Adapted | 5.44 | 13.41 |
| S9 | Adapted | 5.9 | 13.19 |
| S9 | Adapted | 5.45 | 11.15 |
| S14 | Adapted | 9.11 | 18.19 |
| S14 | Adapted | 9.74 | 19.28 |
| S14 | Adapted | 10.06 | 20.34 |
| MC14 | Adapted | 9.71 | - |
| MC14 | Adapted | 7.24 | - |
| MC14 | Adapted | 8.9 | - |
| MF4 | Adapted | 6.01 | - |
| MF4 | Adapted | 5.97 | - |
| MF4 | Adapted | 4.47 | - |
| K3 | Adapted | 3.52 | 25.02 |
| K3 | Adapted | 4.5 | 22.96 |
| K3 | Adapted | 4.41 | 25.95 |
| BH3 | Nonadapted | 5.21 | 29.15 |
| BH3 | Nonadapted | 4.78 | 26.58 |
| BH3 | Nonadapted | 5.11 | 26.61 |
| BH15 | Nonadapted | 4.34 | 13.07 |
| BH15 | Nonadapted | 4.6 | 13.03 |
| BH15 | Nonadapted | 4.36 | 11.74 |
| BR1 | Nonadapted | 6.58 | 10.08 |
| BR1 | Nonadapted | 7.58 | 10.96 |
| BR1 | Nonadapted | 7.84 | 11.29 |
| BR16 | Nonadapted | 7.95 | 13.86 |
| BR16 | Nonadapted | 7.9 | 15.84 |
| BR16 | Nonadapted | 6.82 | 12.98 |
| J3 | Nonadapted | 3.21 | 14.81 |
| J3 | Nonadapted | 3.21 | 14.46 |
| J3 | Nonadapted | 3.39 | 13.83 |
| BU4 | Nonadapted | 1.41 | 9.66 |
| BU4 | Nonadapted | 1.44 | 11.11 |
| BU4 | Nonadapted | 1.58 | 11.54 |
| F10 | Nonadapted | 3.12 | 37.03 |
| F10 | Nonadapted | 2.82 | 39.92 |
| F10 | Nonadapted | 2.57 | 35.47 |
| BH14 | Nonadapted | - | 30.19 |
| BH14 | Nonadapted | - | 30.62 |
| BH14 | Nonadapted | - | 26.64 |
| BR1 | Nonadapted | 5.14 | - |
| BR1 | Nonadapted | 3.53 | - |
| BR1 | Nonadapted | 6.16 | - |
| G11 | Nonadapted | 1.6 | 11.22 |
| G11 | Nonadapted | 1.47 | 10.93 |
| G11 |  | 1.38 | 11.5 |
